# Excitability as a Design Principle in the Immune System

**DOI:** 10.1101/2025.08.31.673412

**Authors:** Yael Lebel, Uri Alon

## Abstract

Growing datasets and mechanistic detail in immunology have outpaced the development of unifying concepts. Such concepts are required to explain the primary goals of immune circuits - strong response to pathogens, tolerance to self, and prevention of collateral damage. A principle that achieves these goals across diverse immune circuits could unify our understanding of the immune system.

Here, we propose that excitability, a concept from dynamical systems, serves this role. We screen thousands of circuits to identify those that generate excitable dynamics, and find a single robust design. We scan the human immune network to find this circuit architecture in dozens of innate and adaptive subsystems.

We provide evidence for excitability in data on longitudinal responses to SARS-CoV-2. Similar motifs underlie T cell activation, autoimmune flares, and tumor immune responses. This conserved motif provides therapeutic targets and suggests that excitability is a core design principle of immunity, bridging molecular and cellular levels.

## Introduction

The innate and adaptive immune systems work by complex networks of communication between effector cells and regulatory cells, using a wide array of cytokines. One way to understand their function is to attempt to isolate circuits of interacting cytokines and cells with specific functions ^1–6^, or to look at the immune system as a network of interacting components^7,8^. If we can find recurring circuit designs with a shared functional principle, we can unify different parts of the immune system in terms of understanding.

The immune system can be considered to have design requirements that apply to many of its subsystems. Three such requirements are strength^9^, noise resistance^10^, and safety^11^. Strength means that the immune system must respond strongly and decisively to threats. Safety means that it must then shut off strongly to avoid collateral damage to healthy tissues. Noise resistance means that weak inputs generated by self-tissue or innocuous stimuli should not trigger a response. An additional design feature - long-term memory - is not considered here. These design goals suggest the need for immune responses that are relevant, decisive, and self-limiting.

One way to obtain all three requirements is by an excitable system. An excitable system has effector and inhibitory components, and produces a strong effector response once a threshold is crossed. Due to the internal dynamics of the effector and inhibitory components, the pulse will shut down even in the continued presence of input. The threshold effect provides noise resistance to the system, ensuring that weak or transient inputs do not result in a potentially harmful response; the strong pulse provides strength to fight threats, such as pathogens or cancer; and the shutdown by the inhibitory component ensures the turn off of the pulse, providing safety from prolonged response that might be harmful to the host.

Excitable dynamics are suggested by certain features of the immune response. Many innate and adaptive responses appear as a strong pulse that then subsides, such as interferon response to virus ^12–14^, inflammatory responses to injury ^15^, or adaptive T cell responses to pathogens ^16^. Effectors such as T cells have a threshold for activation, set in part by antigen-presenting cells such as dendritic cells^17^. Some autoimmune disorders show flare-ups, such as relapsing-remitting Multiple Sclerosis^18^. These flares subside despite the continued presence of self antigen. Excitable dynamics have been studied mathematically in the context of autoimmune flares^18^ and adaptive responses to cancer ^19,20^. However, a systematic analysis of excitability as a potential recurring motif in immunology is lacking.

Excitable systems have been studied in engineering and applied mathematics ^21–24^. A well-known example in biology is the Hodgkin-Huxley equations for action potentials in neurons ^25^ and their subsequent adaptations by FitzHugh^26^, Nagumo^27^, and others ^28,29^. Studies indicate that there are multiple ways to achieve excitability ^30^.

Here we enumerate and compare designs that offer excitability in biological systems. We explore which cytokine and cell-cell interaction circuits can robustly generate excitable dynamics and whether such motifs are found in the human immune network. To do so, we numerically scan a large class of possible circuits of two interacting cytokines or cell types and analyze which architectures produce excitable dynamics. Among thousands of candidates, we find a small set of excitable circuits and identify one optimal design using Pareto optimality principles. This design involves an auto-activating effector that activates its own inhibitor.

We then ask whether this optimal circuit appears in the immune system by searching the Immunoglobe database of cytokine and cell-type interactions. We find that the optimal cytokine and cell-type circuits both recur dozens of times in diverse immune contexts. They are thus a core network motif that appears in different systems and covers about 50% of the immune network. We compare the dynamics of his motif to published data on innate response dynamics in healthy participants inoculated with SARS-CoV-2, to find evidence for excitable dynamics and a pulse rise and deactivation that is insensitive to the timing of viral load dynamics. We discuss instances of the excitable circuit in adaptive immunity, autoimmune diseases and the tumor microenvironment. Finally, we use the excitable circuit model to define targets to reduce or enhance immune responses for therapy.

## Results

### Excitable circuits provide strength, noise resistance, and safety to immune responses

Three desirable features for immune subsystems - strength, noise robustness, and safety - are provided by an *excitable* design (Fig. 1a). An excitable system is defined as having a single stable fixed point such that small stimuli cause a mild response that relaxes back to the fixed point. However, beyond a threshold stimulus, the system shows a very large response (Fig. 1a-c)^31,32^. The system then turns itself off (returns to the stable fixed point) even if the stimulus is still present. It typically has a refractory period until a new pulse can be initiated.

**Figure 1:**
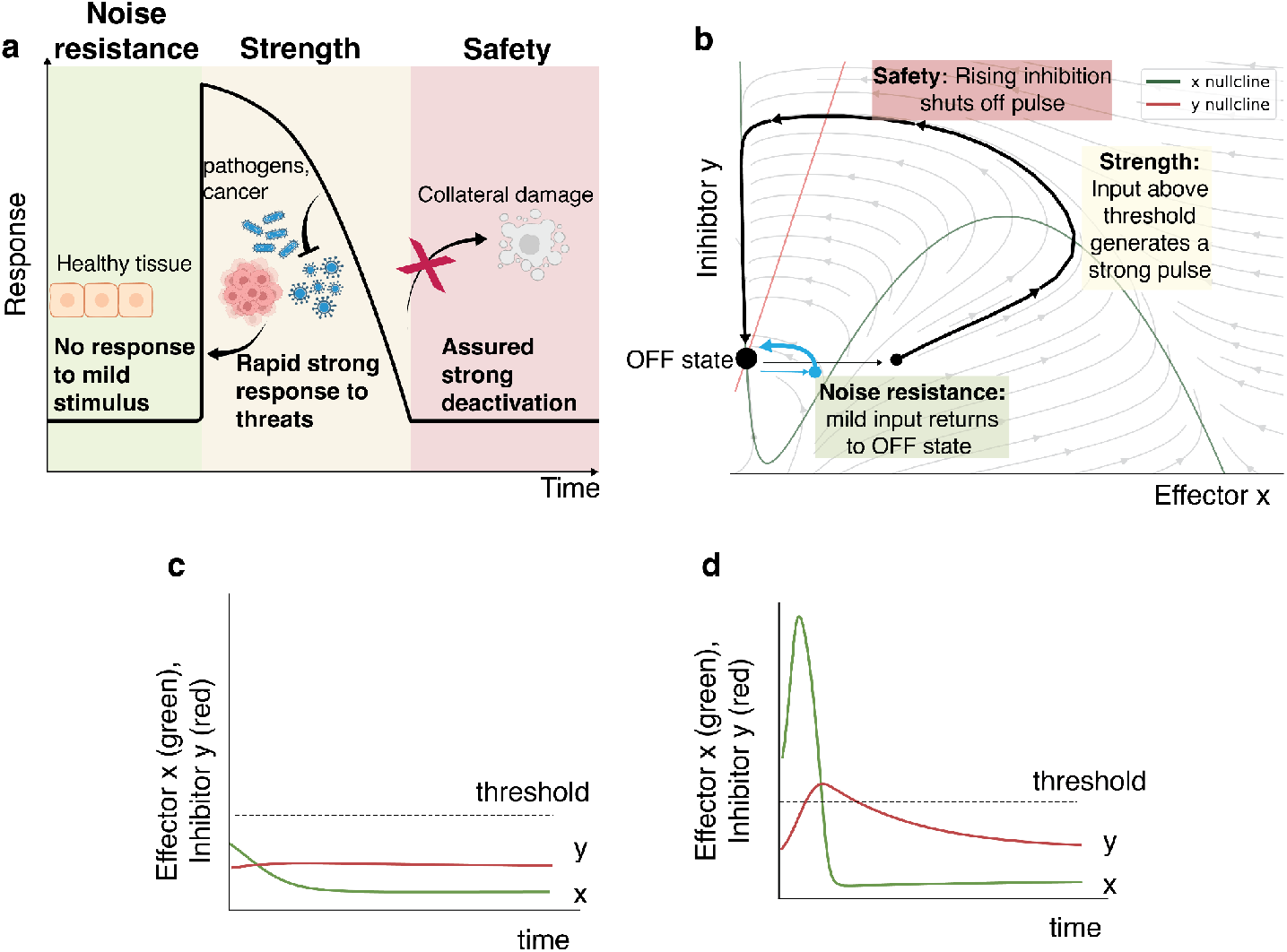
An excitable design can offer strength, noise robustness, and safety to immune responses. a) Three *a priori* specifications for an immune response: noise-robustness - no large response to subthreshold stimuli; strength-rapid and decisive response to supra-threshold stimuli to fight pathogens, cancer, and other threats; safety-assured strong deactivation to avoid collateral damage. b) An excitable circuit can offer these features. It has a single stable fixed point (‘OFF state’) so that weak stimuli converge to the steady state (blue trajectory). Above a threshold input (black trajectory), the effector x rises strongly (strong response). Then, the inhibitor y rises to shut off the response pulse (safety). Above a threshold, dynamics shows a large excursion in phase space before returning to the fixed point. Phase portrait and example trajectories of x and y. c) A suprathreshold stimulus (modeled as an initial condition of x above threshold) causes a pulse, whereas a subthreshold stimulus (d) leads to a mild response that decays to the fixed point.

Robustness to noise is achieved by requiring a stimulus to cross a threshold. This threshold is tuned such that self-tissue signals and random fluctuations are very unlikely to cross it.

The strength of a response is guaranteed by producing a large effector pulse regardless of stimulus strength (provided the stimulus crosses the threshold). Safety is enhanced during a pulse via locality and specificity. Immune cells localize near the point of damage, and immune cells specific to the pathogen are preferably activated.

The decline of the pulse back to the stable fixed point offers safety by preventing excessive effector function. Excitable dynamics effectively cap the amount of collateral damage to the host.

Excitable systems can be implemented in different ways that modify these features quantitatively - certain excitable systems present an all-or-none response, meaning that any suprathreshold input results in nearly the same, stereotyped pulse, whereas other designs have a more gradual sensitivity function, and their response to different suprathreshold stimuli varies in terms of amplitude and duration ^30^.

The circuit equation in b,c,d is 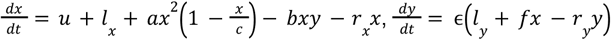 with parameters *l*_*x*_ = 0. 3, *a* = *b* = *r*_*x*_ = *f* = *r*_*y*_ = 1, *c* = 10, *l*_*y*_ = 0. 5, ϵ = 0. 1. The input to the circuit is u. The systems discussed in this study do not assume an explicit separation of timescales between variables, but we use such separation (ϵ ≈ 0. 1) for visual clarity.

### A systematic scan of thousands of cytokine-cytokine circuits yields 32 excitable architectures

Given the potential relevance of excitable circuits to immune responses, we asked how many different ways there are to build an excitable circuit using immune components. We begin by asking which cytokine-interaction circuit designs can provide excitability and whether they are found in the human immune system. Cytokine circuits describe the effect of one cytokine level in the serum on another cytokine’s level. This effect is mediated by cells, for example, by inducing secretion of a certain cytokine, thus increasing its level in the serum, or by deactivating secretion, thus decreasing the cytokine level in the serum.

To search for excitability, we use the definition of excitable dynamics as follows: (i) a single globally stable fixed point at low activity, (ii) initial conditions within a region around the fixed point converge monotonically to it, but beyond this region, the flow is away from the fixed point and then returns to it ^31,32^. These conditions can be tested analytically using the circuit nullclines (Methods).

A single cytokine on its own cannot provide excitable dynamics. The reason is that a one-dimensional dynamical system dx/dt=f(x) cannot provide a pulse for constant input, otherwise there would be concentrations x* that would lead to both a rise f(x*)>0 and a decline f(x*)<0, which is not possible^32^. Excitable models of neuronal action potential can exist on a one-dimensional ring ^29,33^, but these are not readily translated to cytokine concentrations.

We therefore explored which circuits of two cytokines can produce excitable dynamics. In our models, the variables x and y represent the concentrations of cytokines. Cytokines x and y can induce or inhibit their own production and each other’s production. To model cytokine interactions, we use Hill-type production and inhibition terms with Hill coefficients of 0, 1, or 2 (Fig. 2a). This yields a total of 3583 possible circuits.

**Fig. 2.**
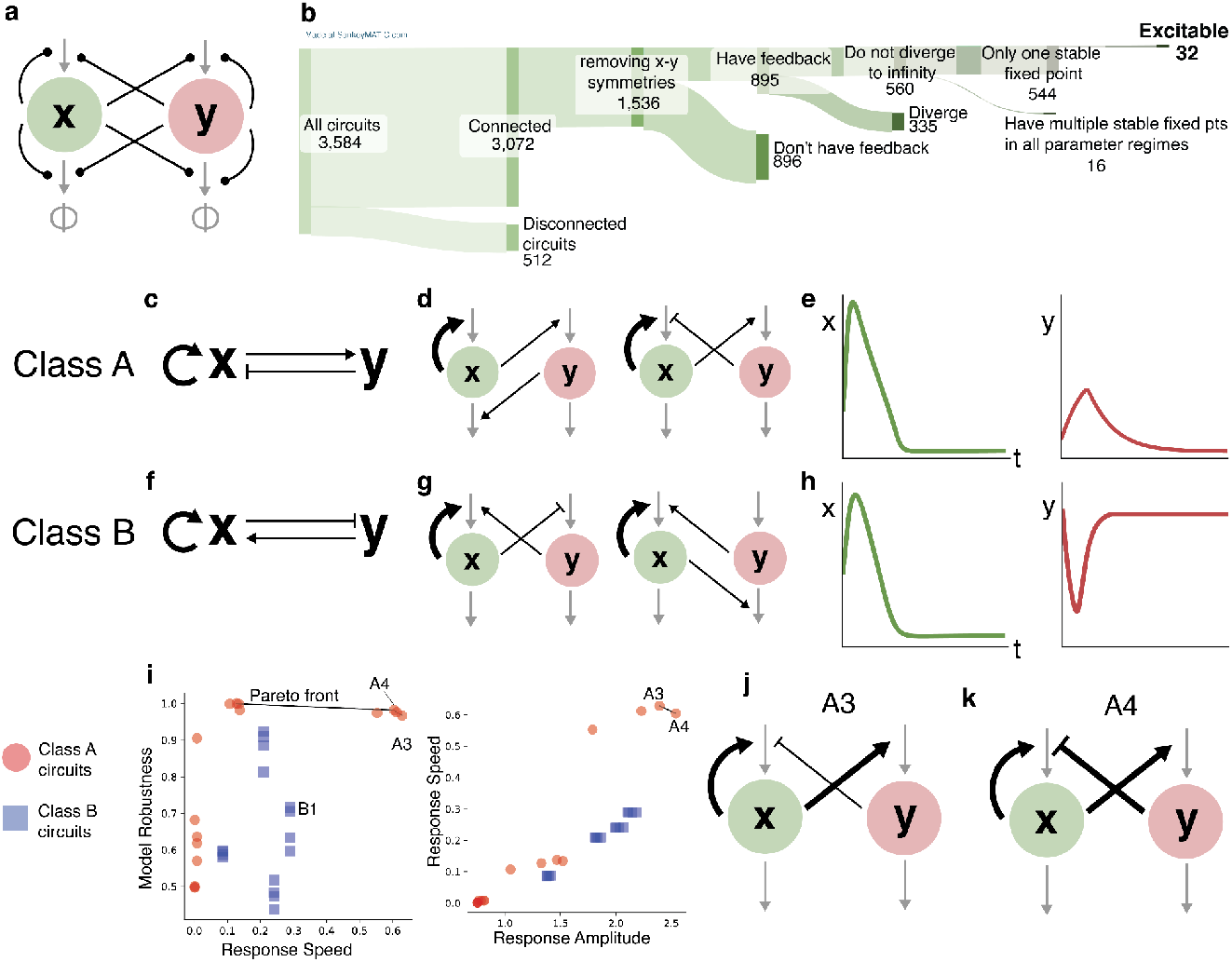
Systematic scan identifies a single excitable cytokine circuit that is Pareto optimal. A. General circuit architecture. Each black line between nodes can represent positive or negative action, with or without cooperativity (Hill coefficient n=2 or n=1, respectively). B. Numbers of circuits with different properties - connected circuits, circuits where the dynamics do not get stuck at the carrying capacity, circuits after symmetry is removed, circuits with no stable ‘ON’ state, excitable circuits. C. Class A excitable circuits: the effector x has a cooperative positive feedback loop and induces its inhibitor y. D. Two examples of the class A circuits. E. X and y dynamics of class A circuits following a suprathreshold input: x and y levels rise and then fall. F. Class B excitable circuits: the effector x has a cooperative positive feedback loop and inhibits its inducer y. G. Two examples of the class B circuits. H. X and y dynamics of class B circuits following a suprathreshold input: x levels rise and then fall, y levels fall and then rise. I. Pareto optimality of circuits A3 and A4 in terms of response strength, speed, and robustness to change in parameters (methods) J. The Pareto optimal circuits A3 and A4.

We find that 32 of these circuits show excitable dynamics (Fig. 2b, methods). The excitable circuits belong to two architectural classes, each with 16 excitable circuits. In the first class, class A, cytokine x induces itself and its inhibitory cytokine y (Fig. 2c). The 16 subtypes differ in whether production or removal is affected and the cooperativity. Two examples of excitable circuits in this class are shown in Fig. 2d.

The second class, class B, is identical except that the cross-regulatory signs are switched - x inhibits the production of its activator y (Fig. 2f). Two examples of excitable circuits in this class are shown in Fig. 2g. The full list of circuit topologies and equations can be found in the SI (Supplementary table 1, supplementary figure 2).

The two classes differ in the dynamics of the inhibitor. In class A circuits, the inhibitor rises then falls during a pulse (Fig. 2e), whereas in class B it falls and then rises (Fig. 2h). Class A has a default (state without stimulus) of no inhibitor, whereas class B has a default of high inhibitor.

### One circuit class is optimal in terms of response speed, strength, and robustness

To compare the 32 circuits, we used a Pareto optimality approach: we defined several performance metrics and asked which circuits outperformed others. We evaluated each circuit in terms of response strength, response speed, and robustness to parameter variation (methods).

To compare the circuits on equal footing, we set as many internal parameters as possible to be the same between the circuits - a mathematically controlled comparison approach ^34–36^. The circuits had, for example, the same threshold for activation and thus identical noise resistance. We then assessed the performance of each circuit across these metrics. We sought circuits that cannot be outperformed across all metrics simultaneously and thus represent Pareto optimal trade-offs between competing features^37^.

We find that class A circuits generally outperform class B circuits in all metrics. Two class A circuits outperformed the others in all tasks (Fig. 2i). In both circuits, cytokine x has cooperative positive feedback on its own production and induces its inhibitory cytokine y cooperatively. Circuit A4 has a cooperative inhibition of x, whereas circuit A3 does not (Fig. 2j,k).

In both circuits an input that crosses the threshold causes a pulse in the x level, followed by a pulse in y levels. There is a refractory period when y levels are high, in which another input to x does not result in a new pulse. Circuit A3 has a shorter refractory period than circuit A4 (SI figure 3).

**Figure 3.**
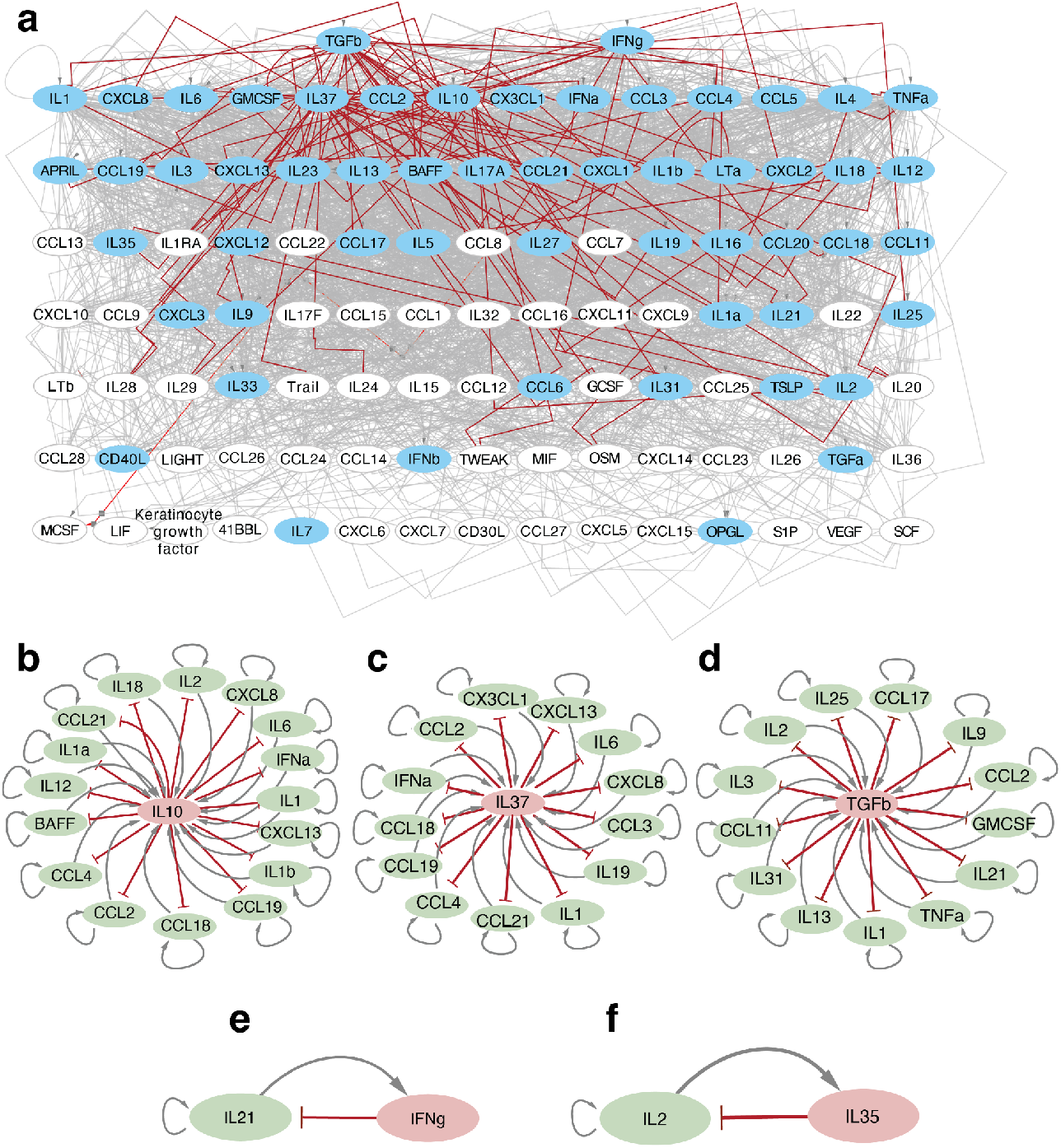
The excitable cytokine circuit is a network motif, with only a few cytokines acting as regulators. a. Human immune cytokine interaction network from Immunoglobe. Cytokines are ordered hierarchically by total degree. Grey arrows represent positive directed regulation between two cytokines, and red arrows represent negative regulation. White nodes are nodes that act as inputs or outputs of the system (in-degree or out-degree = 0). Out of all the blue nodes, which all have both incoming and outgoing edges, all of them participate in an excitable circuit instance. b. b-e) Instances of the excitable cytokine circuit. In each instance, the cytokine in the red ellipse acts as the regulator, and the cytokines in green ellipses act as effectors. Grey arrows represent positive directed regulation between two cytokines, and red arrows represent negative regulation.

### The optimal excitable circuit class is found in 46 cytokine pairs in the human immune system

To explore whether this circuit class appears in the human immune system, we analyzed the network of cytokine interactions from the Immunoglobe database^38^ (methods). The immune system is represented as a graph where each node is a cytokine, and each interaction arrow between x and y indicates that x activates or inhibits the production of y (the arrow also includes which cell types are involved) (Fig. 3a).

In the network, we found all instances of class A circuits defined as a cytokine that enhances its own proliferation and that of another cytokine that inhibits it. We note that in this network, it was not possible to distinguish between the variations within this class.

We find that this circuit class is implemented in 46 different cytokine pairs (Fig. 3b-f). It occurs more often than expected in randomized degree-preserving networks, p=0.0005 (in random networks, there are 30±5 occurrences).

All of the cytokines that are not only input or only output of the network (meaning, their in- and out-degree are >0) participate in an instance of this circuit class. In this sense, the circuit class is found universally in the immune system. We note that this does not mean that all immune systems have excitability, because for some parameter values, these circuits can have different, non-excitable behaviors (see discussion).

Many of these circuits have IL10 as the inhibitor, with activators including IFN-α, IL1a and b, IL6, and TNF-α. Others have TGF-β as an inhibitor. Another major inhibitor is IL37, secreted mainly by monocytes, macrophages, and DCs, with activators including chemokines like CCL2 and CXCL8, as well as cytokines like IL1 and IL6.

We also searched for circuits in class B in which x inhibits its activator y (Fig. 2f). We found 35 examples, a number that is not significant compared to randomized networks (p=0.1, 25±10 in randomized networks).

### Excitable circuit agrees with cytokine response pulse to SARS-CoV-2

Several of the occurrences of the excitable circuit involve pro-inflammatory cytokines and the anti-inflammatory cytokine IL-10. This is consistent with the immune response observed in viral infections.

To test this, we compared the circuit dynamics to published data on the immune response to SARS-CoV-2 in 18 healthy adults^39^, with many cytokines measured daily for 14 days. We defined a composite pro-inflammatory signal using cytokines involved in the class A motif, defined as the rank-sum of the cytokines IFN-α2a, IL-1α, IL-1β, TNF-α, IL-6, CCL4, and CCL2, as the effector node, and IL-10 as the inhibitor.

We compared the measured dynamics to the model-predicted trajectories in the excitable circuit, using Bayesian inference of the parameters (Fig. 4a). The excitable circuit provided good fits to the data in the majority of patients (weighted R^2^=0.64-0.97 (SI figure 4 and SI table 3).

**Figure 4:**
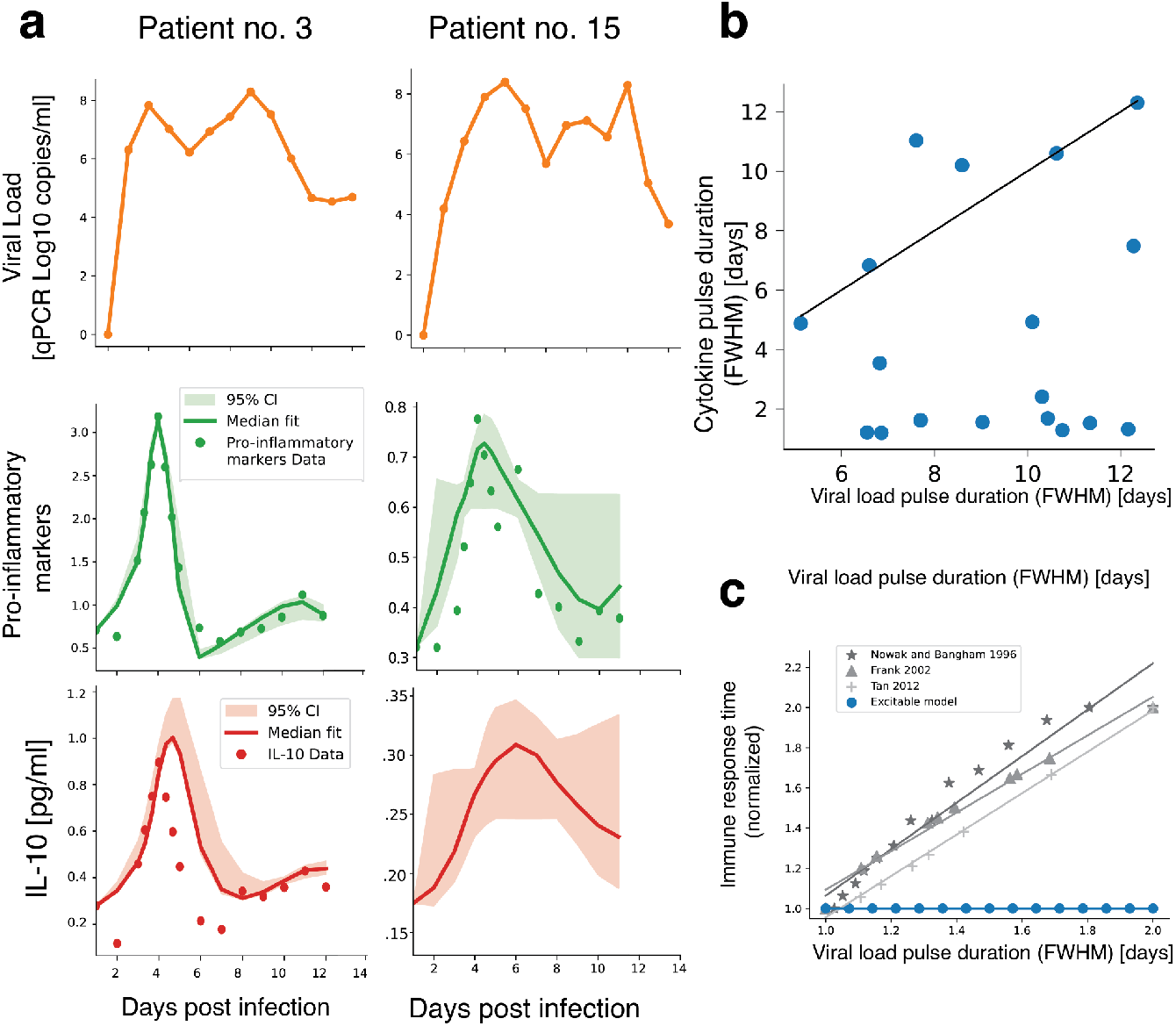
Effector cytokines and IL-10 pulse dynamics agree with an excitable system, and their termination occurs while viral load levels are still high. a) Pro-inflammatory markers (green), serum IL-10 (red), and viral load (orange) in 2 representative patients. Pro-inflammatory pulse ended while viral load was still high. (Trajectories for all 18 patients are in supplementary figure 4) b) Pro-inflammatory pulse ends before viral load. In black: x=y line. Pulse duration (both cytokine and viral load) was calculated using FWHM (Methods). c) The pulse duration is not sensitive to the duration of the input pulse in an excitable circuit (blue); duration is proportional to pathogen duration in other non-excitable models of the immune response. Durations are normalized to the briefest duration. Models are: the 4-equation model of T cell response to a viral infection by Nowak and Bangham 1996 (ref ^12^) (stars); the 3-equation model of response to a viral/parasitic infection by Frank 2002 (ref ^13^) (triangles); and the 3-equation model of the type I interferon response to viral infection by Tan et al, 2012 (ref ^40^) (crosses).

Two of the 18 patients could not be fit by the model (weighted R^2^<0.5), possibly due to measurement noise or atypical dynamics.

### Cytokine response to SARS-CoV-2 terminates even though the viral load is still high, as predicted by the excitable circuit

The excitable circuit terminates its dynamics even though the input is still present. We explored whether the cytokine dynamics in the SARS-CoV-2 dataset agrees with this prediction. As context, we note that several non-excitable mathematical models of immune response show the opposite- a response that is proportional to the duration of pathogen presence. We show three typical examples in Fig. 4c: a 4-equation model of T cell response to a viral infection (Nowak and Bangham 1996, ref ^12^), a 3-equation model of response to a viral/parasitic infection (Frank 2002, ref ^13^), and a 3-equation model of the type I interferon response to viral infection (Tan 2012, ref ^40^). The excitable circuit, by contrast, has a pulse duration that is independent of the pathogen dynamics after a threshold is crossed.

We find that the cytokine effector pulse lasted about a week and was shorter than the period of high viral load, which lasted 12 days on average and varied between individuals (Fig. 4b). To test the relationship between the viral load dynamics and the interferon pulse, we computed the correlation of pulse duration and viral load duration (full-width half max) and found no correlation (r=0.12,p=0.63).

Similar independence of effectors such as type I interferon pulses on viral load can be seen in data of other infections, including influenza and RSV (SI Fig. 1, data adapted from ^41,42^). Qualitative evidence for this type of independent interferon pulse can be found in the innate response to different strains of hepatitis virus ^43,44^.

### A systematic scan of cell-cell circuits identifies a single excitable circuit architecture

Excitability is not limited to the dynamics of cytokines but can also occur in the dynamics of cell populations. This was demonstrated in our recent study of an excitable circuit of effector T cells and Tregs in the context of multiple sclerosis ^18^.

To systematically scan a broad class of cell-cell circuits, we defined a set of biologically plausible equations based on extensions of the model of T cell-Treg interactions ^18^. In this class of circuits, the growth rates of the two cell types, X and Y, are affected polynomially by X and Y, with a carrying capacity term on proliferation that ensures that cell populations do not diverge^45^.

We find that to produce excitability, one needs a minimum of 5 non-zero interactions (monomials). We thus scan all 5-interaction circuits. We also find that one needs a degree of at least 3 in one of the polynomials, and hence we scan all degrees of 0,1,2, or 3 for each monomial interaction (Methods). This defined 6210 possible circuits (Fig. 5a). Circuits with more than five interactions can also provide excitability, but we apply a principle of simplicity (minimal number of interactions) in order to enhance understandability ^37^.

**Figure 5.**
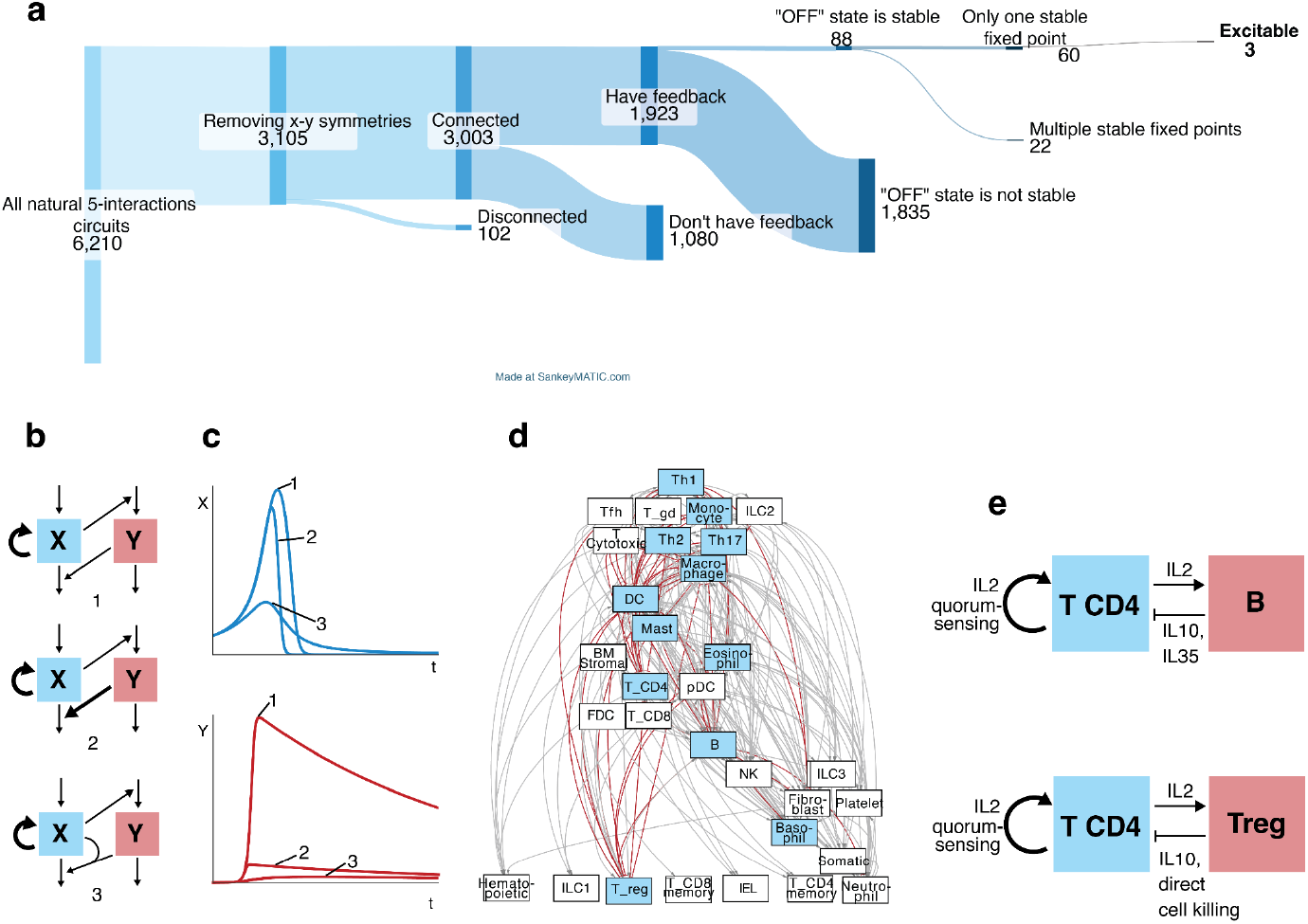
Systematic scan identifies a single excitable cell-circuit architecture. a) Only three cell-cell circuits out of thousands show excitable dynamics, sharing one common architecture. b) The excitable circuits, where bold arrows indicate cooperativity (second-degree polynomial). The equations of the three circuits can be found in the methods. c) Pulse dynamics of the three circuits. d) Cell-cell immune network where nodes are cell types and edges are cytokine interactions, activating (grey) or inhibitory (red). Nodes that participate in the excitable motif are in blue. e) Two instances of cell-cell circuits with the excitable architecture, cytokines are indicated.

We find that out of the 6210 possible circuits, only 3 provide excitability (Fig. 5a,b) (methods). These three share the same general architecture: X enhances its own proliferation rate and activates the production of its inhibitor cell type Y. The three circuits differ in the cooperativity of the inhibitory interaction.

The three circuits respond to an initial input in X with a strong, nonlinear rise of X, followed by a rise in Y that shuts both cell types down. There is a refractory period during which the inhibitor Y is elevated before returning to baseline levels of X and Y (Fig. 5c).

### The excitable cell-cell circuit occurs in 31 cell pairs in the human immune system

To explore whether these circuits appear in the human immune system, we analyzed the network of cell-cell interactions from the Immunoglobe database (Fig. 5d) (methods). In this case, each node is a cell type, and each interaction arrow is a cytokine or set of cytokines sent from one cell type and received by the other. We used igraph’s implementation of the LAD algorithm to find all instances of the excitable circuit (X supports its own and its inhibitor cell type Y growth) (methods). We note here that it was not possible to distinguish between the three variations of the excitable architecture (Fig. 5b) in this network.

We find 31 cell pairs that have the excitable architecture (Table 1). Comparison to randomized networks showed a non-significant trend (p=0.2, 28±4 in randomized networks).

**Table 1.**
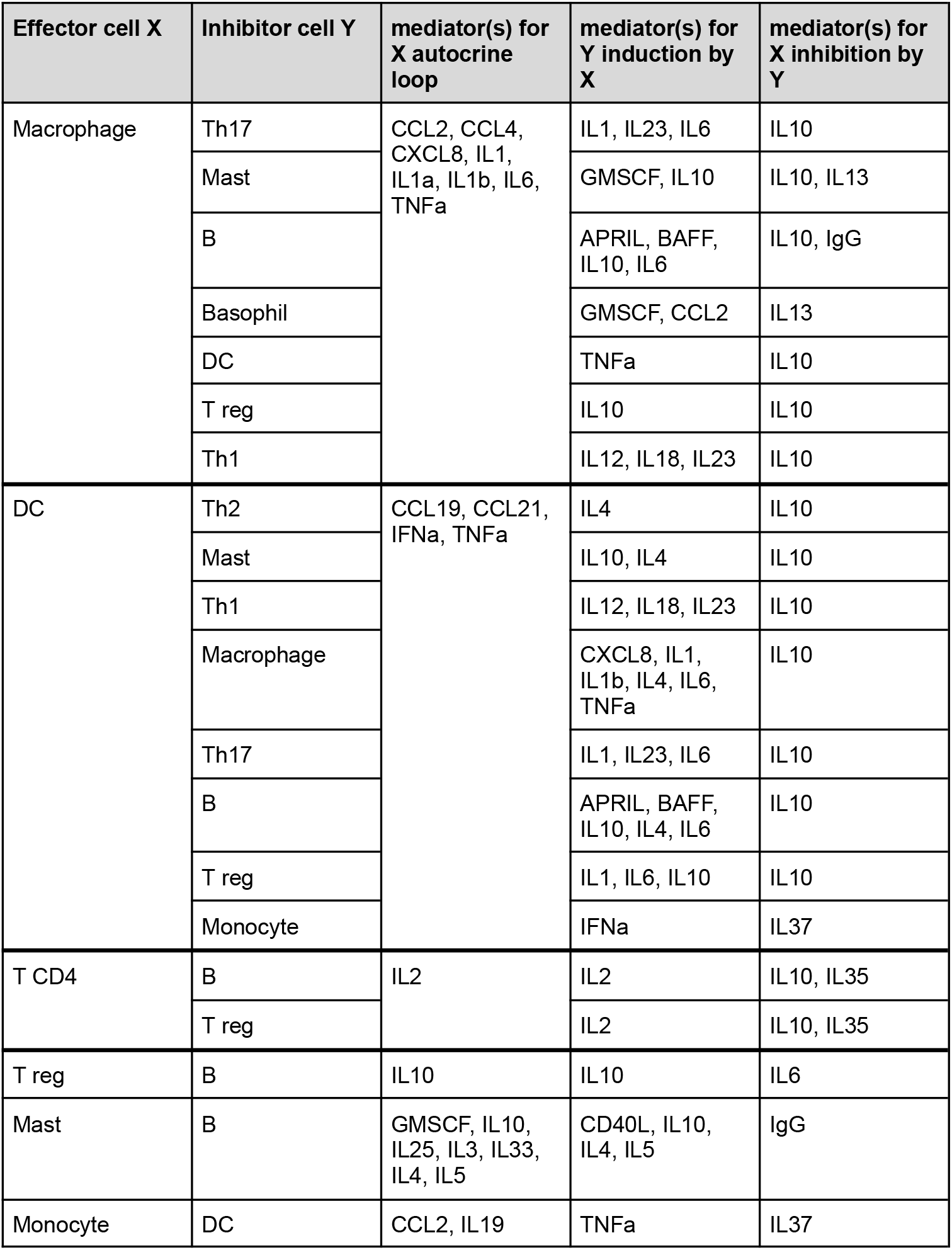
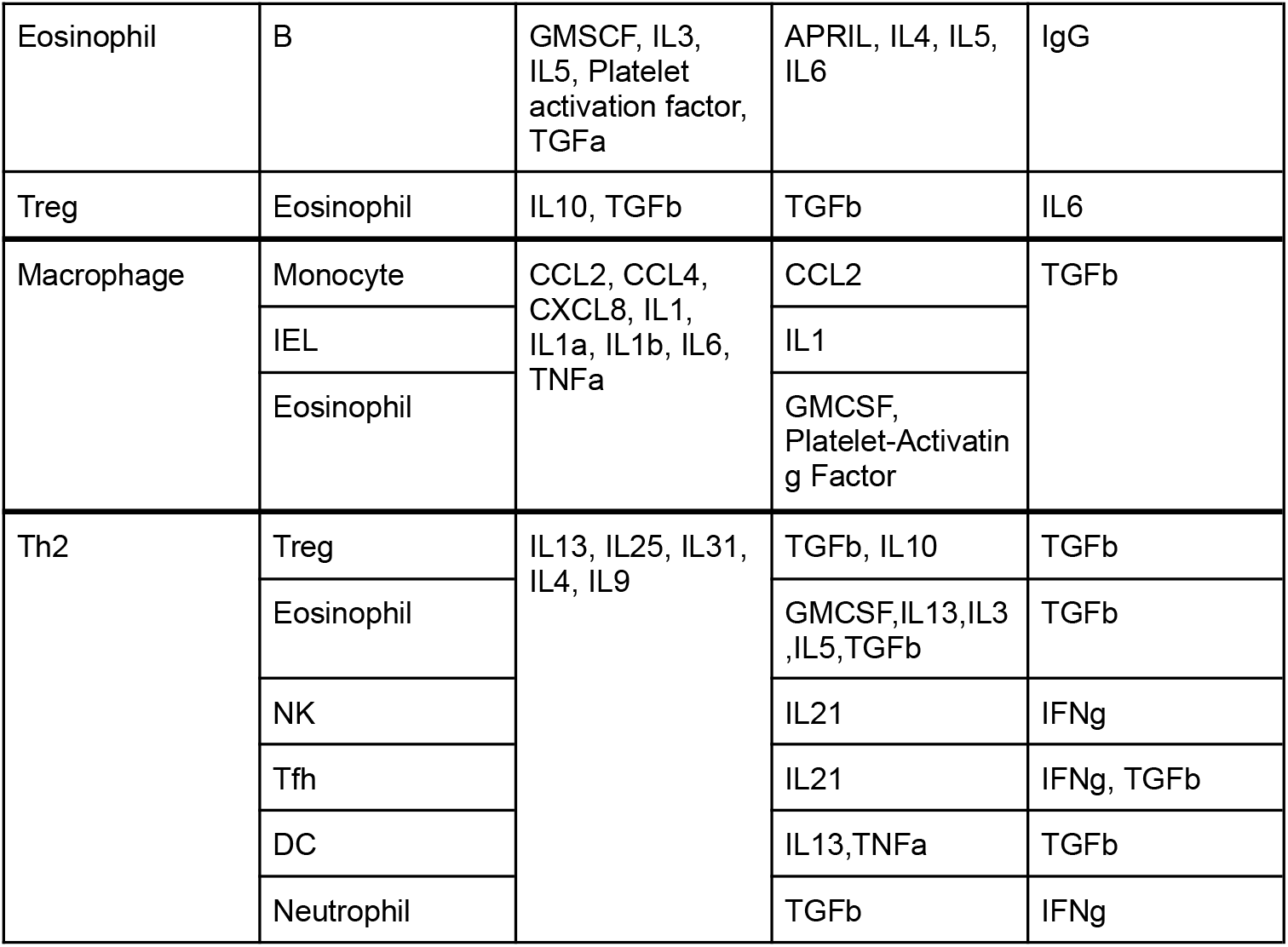
Cell pairs with excitable architecture and the cytokines that participate in the circuits.

| Effector cell X | Inhibitor cell Y | mediator(s) for X autocrine loop | mediator(s) for Y induction by X | mediator(s) for X inhibition by Y |
| --- | --- | --- | --- | --- |
| Macrophage | Th17 | CCL2, CCL4, CXCL8, IL1, IL1a, IL1b, IL6, TNFa | IL1, IL23, IL6 | IL10 |
|  | Mast |  | GMSCF, IL10 | IL10, IL13 |
|  | B |  | APRIL, BAFF, IL10, IL6 | IL10, IgG |
|  | Basophil |  | GMSCF, CCL2 | IL13 |
|  | DC |  | TNFa | IL10 |
|  | T reg |  | IL10 | IL10 |
|  | Th1 |  | IL12, IL18, IL23 | IL10 |
| DC | Th2 | CCL19, CCL21, IFNa, TNFa | IL4 | IL10 |
|  | Mast |  | IL10, IL4 | IL10 |
|  | Th1 |  | IL12, IL18, IL23 | IL10 |
|  | Macrophage |  | CXCL8, IL1, IL1b, IL4, IL6, TNFa | IL10 |
|  | Th17 |  | IL1, IL23, IL6 | IL10 |
|  | B |  | APRIL, BAFF, IL10, IL4, IL6 | IL10 |
|  | T reg |  | IL1, IL6, IL10 | IL10 |
|  | Monocyte |  | IFNa | IL37 |
| T CD4 | B | IL2 | IL2 | IL10, IL35 |
|  | T reg |  | IL2 | IL10, IL35 |
| T reg | B | IL10 | IL10 | IL6 |
| Mast | B | GMSCF, IL10, IL25, IL3, IL33, IL4, IL5 | CD40L, IL10, IL4, IL5 | IgG |
| Monocyte | DC | CCL2, IL19 | TNFa | IL37 |
| Eosinophil | B | GMSCF, IL3, IL5, Platelet activation factor, TGFa | APRIL, IL4, IL5, IL6 | IgG |
| Treg | Eosinophil | IL10, TGFb | TGFb | IL6 |
| Macrophage | Monocyte | CCL2, CCL4, CXCL8, IL1, IL1a, IL1b, IL6, TNFa | CCL2 | TGFb |
|  | IEL |  | IL1 |  |
|  | Eosinophil |  | GMCSF, Platelet-Activating Factor |  |
| Th2 | Treg | IL13, IL25, IL31, IL4, IL9 | TGFb, IL10 | TGFb |
|  | Eosinophil |  | GMCSF, IL13, IL3, IL5, TGFb | TGFb |
|  | NK |  | IL21 | IFNg |
|  | Tfh |  | IL21 | IFNg, TGFb |
|  | DC |  | IL13, TNFa | TGFb |
|  | Neutrophil |  | TGFb | IFNg |

We discuss two examples in more detail (Fig. 5e). The first is a circuit of CD4 T-cells and regulatory T-cells (Tregs) involved in the adaptive response to infection ^46,47^ and in autoimmune disease ^18,48^. In this circuit, CD4 T-cells have an autocatalytic loop via IL2 signaling and quorum sensing^49^; they induce T-reg expansion via IL2; the T-regs eliminate the activity of CD4 T cells via various mechanisms, including cytokines, depletion of IL2, direct and cell killing^50^. This circuit can generate pulses of effector T cell activity that self-terminate through the action of Tregs, independently of remaining antigen, as seen in the context of remitting-relapsing multiple sclerosis, where immune attack stops despite the continued presence of myelin antigen.

Stochastic input (noise) can excite the circuit, and after a refractory period, new pulses can be triggered by subsequent noise (as shown in ref ^18^). Thus, a stochastic input signal can trigger a random series of pulses, with exponentially distributed inter-pulse duration, as observed in relapsing-remitting multiple sclerosis^18^.

A second example is a circuit of CD4 T and B cells. The T-cells support themselves via quorum sensing and IL2 signaling and support B cells via pro-inflammatory cytokine signaling and CD40-CD40L ligands interactions; these cytokines secreted by T cells promote differentiation of B cells towards a regulatory phenotype in which they secrete the antiinflammatory cytokines IL10 and IL35 that inhibit T cells ^51,52^.

Such a circuit was recently identified in the breast cancer tumor microenvironment using one-shot dynamical reconstruction (OSDR)^20^. The OSDR approach allows prediction of cell population dynamics over weeks based on a single biopsy spatial proteomic snapshot by computing the effect of cell neighborhood composition on cell division rate derived from a proliferation marker. OSDR analysis indicated that CD4 T cells enhance the proliferation of neighboring T cells and B cells, whereas B cells inhibit the proliferation of neighboring T and B cells. This results in excitable pulses. In each pulse, CD4 T cells rise first, followed by CD8 T cells.

### Potential drug targets for enhancing or reducing excitable circuit function

We use the cell-cell circuit model to explore potential targets for intervention to reduce the excitable response (useful in the case of autoimmunity) or to enhance it (useful in the case of cancer or hypoimmunity). Analogous results are found for the cytokine circuit.

We define three functional metrics - response amplitude (maximal level of effector x), response threshold (minimal level of x needed to cause a pulse), and refractory period duration (time for inhibitor y to decrease to 1% of its maximum). To find which parameter affects each metric, we perform a sensitivity analysis, defined as the logarithmic derivative of the metric M with respect to the parameter p, 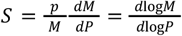 ^53^. Intuitively, sensitivity is the % change in the metric for a 1% change in the parameter.

The model has six parameters (Fig. 6a): autocrine strength *a*, the inhibitory effect *b* of y on x, carrying capacity *c*, x removal rate *d*, the induction effect *e* of y by x, and y turnover rate *e*.

**Fig. 6.**
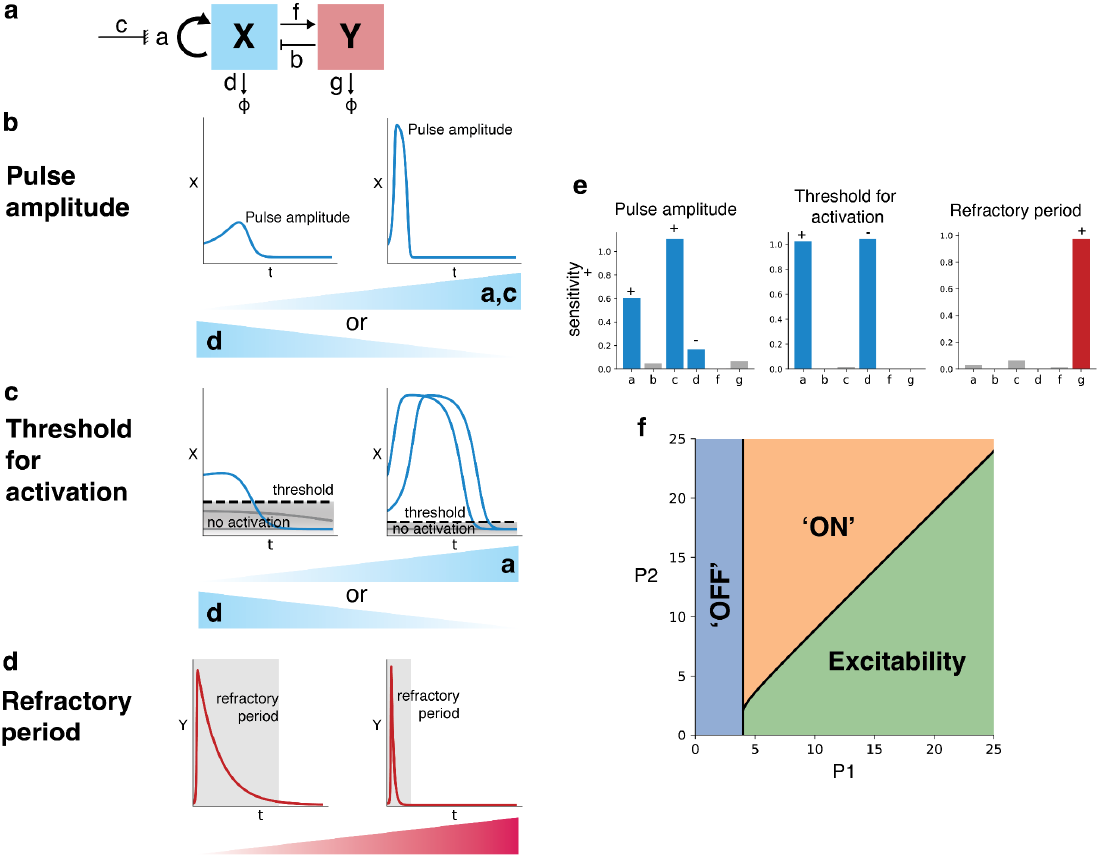
Circuit-to-target analysis suggests targets to enhance or reduce excitable function. **a)** Circuit parameters. b) Increasing X autocrine loop (parameter a) or its carrying capacity c, or decreasing x turnover rate (d), results in stronger pulses; the panels show a=1 (left) and a=5 (right). c) Increasing X autocrine loop (parameter a) or decreasing its removal rate lowers the threshold for activation - the minimal level of X needed to excite a pulse. Three curves are shown for X from three initial conditions, with gray curves below the threshold and blue curves above it. The panels show a=0.1 (left) and a=3 (right). d) Increasing the Y turnover rate (parameter g) shortens the refractory period, defined by the time during which Y is above 1% of the maximum (in grey). The panels show g=1 and g=10. e) Sensitivity of the three metrics to the model parameters, + and - indicate positive or negative effects. f) Phase diagram of the model in terms of dimensionless parameter groups P1, P2 (Methods). Blue - only ‘OFF’ state exists - no activation for any threshold, correlating to hypoimmunity; orange - the ‘ON’ (high effector immunity) is stable, correlating with chronic activation or hyperimmunity; green: excitability as discussed in this paper. Circuit equations are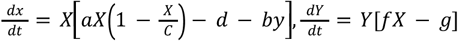 (carrying capacity for Y was omitted for clarity and simplicity of analysis). Sensitivity was calculated around the parameter set *a* = *b* = *c* = *g* = 1, *d* = 2, *c* = 100.

Only some of the parameters have large effects on the metrics (Fig. 6b-e). Response amplitude rises with the autocrine strength *a*, and with carrying capacity, *c*. This is due to the self-amplification effect, which determines the pulse amplitude that reaches near the carrying capacity. Response amplitude decreases with the removal rate of x, *d*.

The threshold for activation rises with the autocrine strength *a*, and decreases with the removal rate of x, *d*. Thus, both amplitude and threshold are not substantially affected by parameters related to the inhibitor y.

The y parameters mainly affect the refractory period duration, which decreases with y removal rate, g. It is not sensitive to parameters related to the activator x. This is because the refractory period is the time when the inhibitor is high and effector is low, a period of time determined by the speed of y removal.

The paracrine interaction parameters f and b are not sensitive targets for these metrics of this circuit.

One can also ask about the range of behaviors of the circuit when parameters are changed by large amounts (Fig. 6f). Typical excitable circuits such as the Hodgkin-Huxley model for neuronal action potentials can show a range of behaviors-stuck in the OFF state, a new stable ON state, bistability between ON and OFF, and oscillations.

The cell circuit studied here has a more limited range of behaviors- it shows excitability for a very large range of parameters (dimensionless parameter groups larger than a threshold value). When parameter groups become very small it can show two ‘pathological’ behaviors - always OFF, and always ON in a new stable fixed point of high activity. These behaviors may mimic pathologies of of hypo- and hyper immunity.The circuit can show no oscillations or bistability, and in this sense is ‘protected’ from oscillatory pathologies.

## Discussion

We suggest that excitability is a recurring design principle in immunology and show that circuits with excitable potential recur throughout the immune system. To explore how many excitable circuits exist, we scanned a large class of two-cytokine and two-cell circuits. Out of thousands of possible circuits, only one basic design is excitable and superior to other designs in terms of strength, speed, and robustness to parameter variation: a cooperatively self-enhancing effector that induces its own inhibitor. We show that this design is a network motif that appears dozens of times in the human immune network with different cells and cytokines. Almost all cytokines participate in an instance of this motif. We reanalyze published data to provide evidence that cytokine response to viral infection has excitatory-like properties, with a pulse that shuts down before viral load diminishes. We discuss excitable circuits in various contexts and use the excitable circuit model to find potential targets to enhance or repress immune response.

An excitable circuit can provide three basic requirements for effective immune function: noise resistance, where a sub-threshold input leads to no response; strength, where a supra-threshold signal leads to a large pulse of activity that is independent of further antigen input; and safety in which the response turns itself off to avoid collateral damage.

We find that many different effector cytokines participate in the excitable design, but the number of inhibitory partners is much more limited. Only four inhibitors, IL10, IL37, TGFβ and IL35, account for almost all inhibitory cytokines. It is interesting to ask why the number of effectors is much larger than the number of inhibitors. One possibility is that the inhibitors play mainly a dynamic role in shutting off the response, whereas each effector needs to perform a cognate function against specific threats. A shared inhibitor can coordinate shutoff between a large set of effectors, providing global shutdown to bespoke multi-effector response pulses.

In addition to circuits of cytokines, we also scanned circuits of two interacting cell types. We find one circuit out of several thousand possible ones that can give rise to an excitable pulse of cells. This excitable circuit has a similar architecture to the cytokine circuit- a cell type that enhances its own proliferation cooperatively and induces its inhibitory cell type. This circuit occurs in 21 different cell type pairs in both innate and adaptive immunity. Examples include a CD4T cell-B cell circuit previously suggested to cause excitable pulses of anti-cancer immune attack^20^, and a CD4 T cell-Treg circuit previously suggested to underlie excitable autoimmune pulses in multiple sclerosis^18^.

It is intriguing that the same essential design (autocatalytic x that activates its inhibitor y) occurs on both levels of organization-cells and cytokines, hinting that the space of excitable systems may be limited to a single design that is biologically feasible and effective.

It is important to note that the widespread occurrence of a circuit that can show excitability does not mean that all immune functions are excitable. Some aspects of immune response seem to have the features of an excitable system, such as the cytokine storm and interferon responses observed in severe COVID-19 cases^54^. A quick response followed by a decline is a very common motif across adaptive and innate immunity^14^. The refractory phase is best known in macrophages^55–58^ and exists in T cells (though less understood mechanistically)^59,60^. A threshold for activation is common but not universal, present in T cells^61^ and mast cells^62,63^ but not macrophages - where some responses are graded in response to input and seem to have no threshold. Other parts of the immune system are not excitatory at all, such as long-term memory of B cells and T cells, which does not shut itself off.

We used the circuit model to search for potential targets to enhance or reduce the excitable function. We find that the autocrine loop, in which the activator enhances its own production, is a sensitive target - it increases both the amplitude and reduces the activation threshold of the circuit. Thus, inhibiting this loop is predicted to make false activation rarer and weaker, which is a desirable outcome in autoimmune diseases. This relates to recent findings in fibrosis, where a circuit model predicted that inhibiting the autocrine loop of myofibroblasts would reduce and prevent fibrosis, a prediction that was validated in mice models of heart and liver fibrosis^64^.

Further targets include the removal rate of the effector x, which also controls amplitude and threshold. The refractory period is mainly affected by the removal rate of the inhibitor y. Interestingly, the paracrine interactions between effector and inhibitor are not sensitive targets for these metrics.

The excitable design has drawbacks and fragilities. If the threat is eliminated early, the pulse may continue its course, causing excess collateral damage ^65^. The excitable circuit can have potentially pathological behaviors if its parameters are changed by a large amount. This can cause bifurcation to undesired states such as a stable fixed point of permanent high activity. Some of these bifurcations may correspond to immunopathologies (Fig 6f).

One limitation of the present study is that it ignores the uniqueness of each immune circuit in terms of biochemistry, cell biology, anatomical distribution, and pathogen context. It takes a generalist approach, a view from 30,000 feet, to highlight principles on the level of circuit architecture. These principles can guide the search for classes of targets for intervention, but detailed biochemical knowledge is required to implement such interventions for each particular cytokine or cell circuit. Another limitation is ignoring spatial aspects-one can imagine immune flares in certain regions of a tumor, for example, and not others.

Excitability is familiar to biologists from the Hodgkin-Huxley (HH) model of neuronal action potential. The HH model is foundational in neurobiology, guiding experimental studies on ion channels and other systems. The HH model also gave rise to a field in applied math based on the Fitzhugh-Nagumo simplification. Refractory periods and stereotyped effector pulses are also found in immunology, not only at the level of cells, but also in entire immune subsystems.

We conclude that excitability is a design principle in the immune system and that one basic circuit design (with some variations) provides excitability across diverse adaptive and immune settings, implemented with different cells and cytokines. This provides a mathematical framework that may unify aspects of immune responses.

## Methods

### 1. Comparison of immune response length to viral infection duration in other models

#### 1.1 Nowak and Bangham 1996

The model equations are:

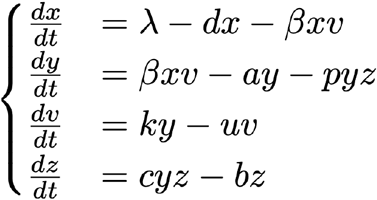

With x the number of uninfected cells, y the number of infected cells, v the free virus particles, and z the CTL immune response. The equations were integrated from t=0 to t=100 in time steps of 0.01 using the Euler method for integration, with initial conditions set to be x=0.99, y=0.01, v = 0.0, z= 0.01, as in the original paper.

The model parameters, taken from the original paper, were:

- λ = 1.0
- d = 0.01
- β = 1.0
- a = 0.5
- p = 1.0
- u = 20.0
- c = 1.0
- b = 0.05
- k ∈ [0.5,2.5]

#### 1.2. Frank 2002

The model equations are:

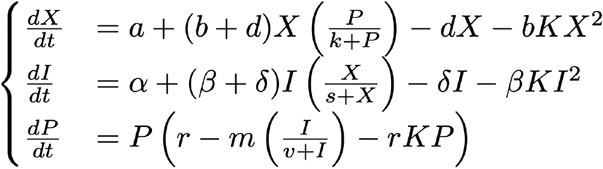

With X concentration of molecule that increases directly in response to parasitic attack, I is the concentration of immune cells that can kill the parasite, and P is the concentration of parasites within the host.

The model parameters, taken from the original paper, were:

- a = 0.05
- b = 2.5
- d = 2.5
- α = 0.05
- β = 2.5
- δ = 2.5
- k = 1e4
- K = 1e-8
- s = 1e4
- r ∈ [1.5, 3]
- m = 5.0
- v = 1e5

The equations were integrated from t=0 to t=40 in time steps of 0.01 using the Euler method for integration. Initial conditions were set to be *X* = *a*/*d*,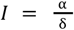, *P* = 1. 0, as in the original paper.

#### 1.3. Tan 2012

The model equations are:

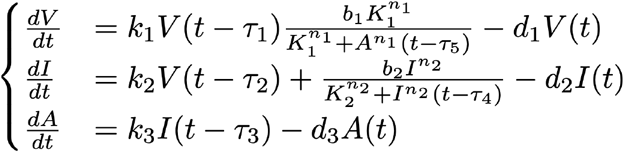

WhereV (mol/L), I (mol/L), and A (mol/L) are the concentrations of viral mRNAs, IFNs, and AVPs, respectively.

The model parameters, taken from the original paper, were:

- k1 = 4.0
- k2 = 0.3
- k3 = 0.1
- d2 = 0.7
- d3 = 0.12
- b1 = 10
- b2 = 80
- K1 = 33
- K2 = 0.1
- n1, n2 = 1
- d1 ∈ [0.035, 0.35]

The delayed differential equations were integrated using ddeint^66^, an open-source package for delayed differential equations based on scipy’s odeint^67^. The equations were integrated from t=0 to t=100 in time steps of 0.1.

#### 1.4 Excitable system

The equations used were:

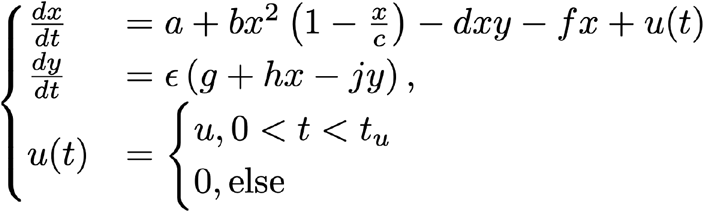

The model parameters were:

- a = 0.3
- b = 1
- c = 100
- d = 1
- f = 1
- ε = 0.1
- g = 0.5
- h = 1
- j = 1
- u = 6
- t_u ∈ [5,25]

The equations were integrated from t=0 to t=50 in time steps of 0.01 using the Euler method for integration. Initial conditions were: x= 0.2, y=0.7.

#### 1.5 Extracting infection and immune response duration

For each of the models, the duration of the infection and the immune response was determined using full-width half max (FWHM), i.e, the duration of time in which each of the variables was above half of its maximum value.

### 2. Analysis of Longitudinal type I interferon and SARS-CoV-2 Patient Data

We analyzed longitudinal clinical measurements of type I interferon (IFN1) and viral load in patients experimentally inoculated with SARS-CoV-2, based on publicly available data from [Ref ^39^]. The dataset included daily measurements of plasma IFN-a2a levels and twice-daily measurements of viral RNA concentration for n=18 individuals, starting from one day before the day of inoculation.

To compare the dynamics of the IFN-a2a pulse and the viral load, we extracted the pulse amplitude (maximum IFN-a2a value) and pulse duration, defined as the full width at half maximum (FWHM) of the IFN-a2a time trace. Viral load amplitude and duration were similarly defined.

To calculate FWHM, we first located the maximum value of the IFN-a2a time series and computed the half-maximum level. We then identified the time points at which the IFN-a2a signal crossed this half-maximum on the rising and falling edges of the pulse.

To achieve sub-sampling accuracy, we used linear interpolation around the crossing points. Specifically:

1. We identified the indices of the first and last values that crossed the half-maximum threshold;
2. For each crossing point, we interpolated between the surrounding two time points using scipy.interpolate.interp1d;
3. The FWHM was calculated as the time difference between the two interpolated crossing times.

To assess whether the IFN-a2a pulse was shaped by viral load dynamics (as would be expected in a pathogen-limited model), we computed:

- The Pearson correlation coefficient between IFN-a2a and viral load amplitudes
- The Pearson correlation between IFN-a2a pulse duration and viral load duration.

We also computed the coefficient of variation (CV) of both IFN-a2a amplitude and viral load across individuals. Statistical significance was evaluated using two-sided p-values for Pearson correlation. All analyses were performed in Python using numpy, scipy.stats, and pandas.

### 3. Systematic Scan of Cytokine-Cytokine Circuits

#### 3.1 Circuit Space and Excitability Scan

To systematically enumerate circuits with exactly three non-zero regulatory interactions, we generated all possible configurations in which the production and removal terms of x and y could include either positive, negative, or null regulation. Each circuit was defined by assigning an interaction value from the set {–2, –1, 1, 2} to four regulatory sites for x and four for y. The interaction values represent different types of regulation: ±1 for non-cooperative and ±2 for cooperative activation or inhibition; zero indicates the absence of interaction. We generated all configurations in which exactly three positions were assigned a non-zero interaction value, with the rest set to zero. We further filtered these combinations to retain only those with specific distributions of non-zero interactions on x and y (e.g., 3:0, 2:1, 1:2, 0:3), ensuring full coverage of all topological possibilities with three total interactions. This process yielded 3583 unique circuits. Each regulatory interaction was modeled using Hill-type activation or inhibition functions with Hill coefficients n=0 (no interaction), 1 (non-cooperative), or 2 (cooperative).

Each circuit was represented by a system of ordinary differential equations (ODEs):

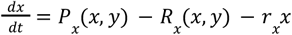

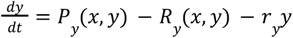

Where *P*_*x*_, *P*_*y*_ (production of x and y, respectively) and *R*_*x*_, *R*_*y*_ (removal of x and y, respectively) are products of Hill functions representing activation and inhibition, and *r*_*x*_, *r*_*y*_ are linear degradation rates.

#### 3.2 Screening for excitable circuits

We first generated all combinatorially possible topologies with 3 interactions, i.e, the number of Hill functions in the product of Px, Py, Rx, Ry was exactly 3. The total number of possible circuits was 3583.

The following screening steps were taken:

1. We first screened out circuits that were not connected, i.e, there was no interaction in either Px or Rx involving y and no interaction in Py or Ry involving x, remaining with 3072 circuits;
2. Removed circuits that are x-y symmetric to other circuits, remaining with 1536 circuits;
3. We then removed circuits that don’t have feedback between x and y and remained only with circuits in which x feeds to y and y feeds back to x, summing up to 640 circuits;
4. We remove circuits that are not contained and diverge to infinity, namely, circuits in which one variable has only positive interactions and no negative interactions with itself or the other variable, and remain with 560 circuits. For the remaining circuits after this stage, we computed the nullclines numerically using Sympy’s symbolic equations solver.
5. Circuits for which a stable fixed point far from the zero was created were eliminated, remaining with 540 circuits. The stability of the fixed point was estimated by calculating the eigenvalues of the system’s Jacobian at the fixed point.
6. We next screened for threshold behavior. Only circuits for which in some parameter range an N-shaped nullcline was created, i.e, one of the nullclines had both a local minimum and a local maximum, were screened for. In this stage, we remained with 32 circuits that were categorized as excitable.

The table of equations for the remaining 32 excitable circuits can be found in the supplementary.

#### 3.3 Mathematically controlled comparison

To compare the circuits using different performance metrics, we followed the principles set by Michael Savageau in his works on mathematically controlled comparison ^34–36^. According to this method, as many internal parameters of the systems in comparison should be set to be the same. Since the equations describing the dynamics of the different circuits are structurally different, we opted to set the topological parameters of the circuits the same across the circuits. Namely:

1. For each model i, the N-shaped nullcline *N*_*i*_(*x*) has a local minimum 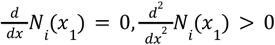 and a local maximum 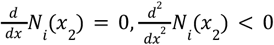 for the same x1, x2 across all models.
2. For each model, the N-shaped nullcline has the same minimum *N*_*i*_ (*x* _1_) and maximum *N*_*i*_ (*x* _2_) = 0 across all models.
3. The steady-state was in the same place, namely the N-shaped nullcline *N*_*i*_ (*x*) and the other nullcline *L*_*i*_ (*x*) crossed at the same *x*^*^ for all models and had the same *N* _*i*_ (*x* ^*^) and *L*_*i*_ (*x*^*^) across all models.

The same input in x was inserted into all models as initial conditions, and their dynamics were evolved using the Euler method.

To compare the circuits, the following performance metrics were defined:

a. Response strength was defined as the maximal x output.
b. Response speed was defined as the maximal x output divided by the time it took to get to this maximum.
c. The refractory period was estimated by the maximal y output.
d. Model robustness was estimated using the following algorithm:
  i. For each model, 1000 parameter sets were drawn from a Gaussian distribution with a standard deviation of 10% around the original value;
  ii. All parameter sets were checked to see if criterion 6 in the circuit can still hold (namely, whether there is a local minimum and a maximum, and whether the nullclines cross at a single stable fixed point).

We identified optimal circuits using a non-dominated sorting approach: a circuit is optimal if no other circuit outperforms it in all metrics simultaneously. This analysis revealed that a single circuit - where cytokine x activates itself and induces its inhibitor y - consistently occupied the Pareto front across all four metrics. This class 1 excitable circuit was thus selected as the optimal design, combining strong, fast, and robust pulse generation with an effective refractory mechanism.

### 4. Network Search for Excitable Cytokine Motifs

#### 4.1 Construction of Cytokine-Cytokine Interaction Network from ImmunoGlobe

The Immunoglobe database [Ref ^38^] provides a rich network of immune interactions, including cell types and cytokines, along with the direction, nature, and biological processes associated with each interaction. To identify dynamical motifs at the cytokine level, we reconstructed an effective cytokine-cytokine interaction network by inferring cell-mediated cytokine interactions from this bipartite immune system graph.

Our network construction followed these steps:

1. Node classification: We first classified all nodes in the Immunoglobe dataset as either cytokines or cells based on their annotated type.
2. Removal of high-connectivity cytokines: Before constructing the interaction network, we removed two cytokines - IFN-γ and TGF-β - from the dataset. These cytokines appear in a large fraction of interactions (28% and 21% of cell types, respectively) and could dominate circuit statistics due to their broad, non-specific regulatory roles.
3. One-step mediated cytokine interactions: We inferred indirect interactions where cytokine x affects cytokine y through a mediating cell:
  - Step 1: Cytokine x acts on a cell (e.g., via receptor binding or signaling);
  - Step 2: That cell, in turn, produces cytokine y in response.
  - These paths were extracted by finding cytokine → cell and then cell → cytokine edges and merging them on the shared intermediate cell.
4. Interaction sign and biological annotation: Each inferred edge between cytokines was assigned a net interaction sign using a conservative rule:
  - If either action in the path (e.g., activation, inhibition, killing) was inhibitory, the net effect was labeled Negative.
  - Otherwise, the interaction was labeled Positive.
5. Final network structure: The final cytokine-cytokine interaction network was a directed, signed graph where each edge included:
  - Source and target cytokines
  - Interaction sign (Positive or Negative),
  - Intermediating cell type (Mediator).

Where multiple mediating paths existed between two cytokines, we aggregated all paths by combining distinct mediators and merging associated immune processes. This yielded a biologically annotated and directionally consistent cytokine interaction network that better reflects functional pathways in immune regulation.

#### 4.2 Motif Search

Using this cytokine-cytokine network, we searched for instances of the excitable circuit motif identified in Section 3. Specifically, we searched for directed two-node motifs with:

- A self-activating cytokine x;
- A positive regulation from x to cytokine y,
- A negative regulation from y back to x.

We used the Labeled Adjacency Descriptor (LAD) algorithm in igraph to identify all such three-node subgraphs with the required directionality and edge labels (Positive, Negative). For each match, we recorded the identities of cytokines x and the cell type/s mediating the interaction (if inferred).

#### 4.3 Statistical Enrichment of Excitable Motifs

To assess whether the observed frequency of excitable motifs in the cytokine-cytokine network exceeds what would be expected by chance, we generated null models by rewiring the network while preserving key structural constraints. Specifically, we constructed randomized networks that:

- Maintained the original node set (cytokines and their labels),
- Preserved the total number of edges,
- Retained the direction and sign (“ Positive” or “ Negative”) of each interaction,
- Avoided duplicate interactions (i.e., no multiple edges of the same action type between the same source and target).

We implemented a custom edge rewiring procedure using a pairwise swap algorithm. For each randomized network, we:

1. Randomly selected two directed edges (*u* → *v, action*1) and (*w* → *x, action*2)
2. Proposed swapping their targets to form(*u* → *x, action*1) and (*w* → *v, action*2) Accepted the swap only if it did not introduce a duplicate edge of the same interaction type.

This procedure was repeated for 10,000 times per graph to ensure complete randomness. The resulting randomized networks preserved the degree distribution, edge sign distribution, and edge uniqueness constraint.

To evaluate whether excitable motifs were enriched in the immune network beyond random chance, we generated 10,000 degree-preserving randomized networks using igraph. These networks maintained the same in- and out-degree distribution across cytokines but randomized edge connections, thereby destroying higher-order motifs while preserving node-level statistics.

We applied the same LAD-based motif search to each randomized network and computed the number of excitable circuit matches. Comparing the observed count of 32 to the null distribution, we computed an empirical p-value:

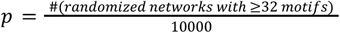

### 5. Systematic Scan of Cell-Cell Circuits

#### 5.1 Circuit Class Construction

To investigate whether excitable dynamics arise in the interaction between immune cell populations, we developed a generalized framework for modeling the dynamics of two interacting cell types, denoted X and Y.

We modeled the temporal dynamics of the cell populations with the following general form:

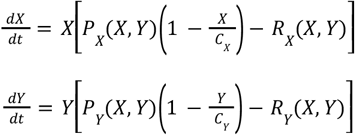

With *P*_*X*_,*P*_*Y*_ representing the production terms of X and Y respectively, *R* _*X*_,*R*_*Y*_ the removal of X and Y respectively, and *C*_*X*_,*C*_*Y*_ the carrying capacities of X and Y respectively. This formulation ensures that when *X* = 0 or *Y* = 0, the corresponding population remains at zero, reflecting the biological irreversibility of extinction in the absence of cell division. This form also ensures that cell populations do not diverge to infinity.

To define the space of possible interaction topologies, we used a polynomial expansion approach. Specifically, we considered that each function *P*_*X*_,*P* _*Y*_, *R*_*X*_,*R*_*Y*_ is a sum of terms of the form:

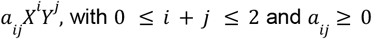

This includes all monomials of total degree up to 2, resulting in an overall degree of 3 (including the carrying capacity). This formalism gives rise to interactions such as:

- Constant background proliferation (free term in *P* _*X*_, *P*_*Y*_) or removal (free term in *R* _*X*_, *R*_*Y*_);
- Linear and quadratic self-activation or self-limitation,
- Cross-activation or inhibition,
- Cooperative inhibition via terms like *XY, X* ^2^ or *Y*^2^.

#### 5.2 Screening for excitable circuit

The overall number of combinatorially possible circuits with 5 interactions is 8568. We first filter to retrieve “ natural” 5 circuit interactions by:

- Removing circuits in which all 5 interactions appear in one variable (either X or Y) with essentially only one of the variables having any dynamics, remaining with 8316 circuits;
- Removing circuits in which there are no removal terms in either X or Y, remaining with 6990 circuits;
- Removing circuits in which there are no production terms in either X or Y, remaining with 6210 circuits.

We next filter the dubbed “ natural” circuits to screen for excitable circuits:

- Removing circuits that are symmetric to existing circuits under X-Y symmetries, remaining with 3105 circuits;
- Removing disconnected circuits, remaining with 3003 circuits;
- Removing circuits that don’t have feedback, remaining with 1923 circuits;
- Removing circuits in which the “ OFF” state in the origin (X=0, Y=0) is unstable or half stable, remaining with 1304 circuits;
- Removing circuits in which there are other stable fixed points, on top of the “ OFF” state at the origin, for all parameter values, remaining with 60 circuits;
- Filtering to get circuits that are excitable for a robust range of parameters, remaining with 3 circuits.

#### 5.3 Equations for the 3 resulting excitable circuits

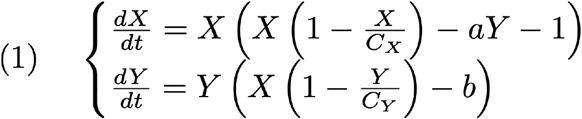

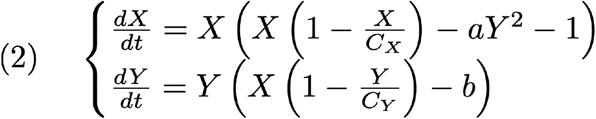

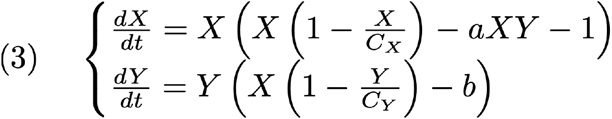

### 6. Network Search for Excitable Cell-Cell Motifs

#### 6.1 Construction of the Cell-Cell Interaction Network

To determine whether excitable cell population circuits appear in real immune systems, we constructed a directed network of cell-cell interactions based on cytokine signaling paths reported in the Immunoglobe database. The original data describe interactions between cells and cytokines, where each edge is annotated with a named action (such as Activate, Secrete, Inhibit, or Kill) and associated immune processes. Our goal was to reconstruct a functional graph of interactions between immune cell types by parsing, filtering, and abstracting these cell-cytokine-cell paths.

We began by parsing interaction edges from ImmunoGlobe’s annotated edge strings, extracting the source (e.g., a cell), the action type (e.g., Activate), and the target (which could be a cytokine or another cell). We excluded entries with ambiguous or indirect actions, such as “ Polarize,” and retained only interactions relevant for population-level effects: Activate, Survive, Secrete, Inhibit, and Kill.

We also filtered out non-immune or non-leukocyte cell types such as keratinocytes, hepatocytes, epithelial and endothelial cells, and tumor models. The resulting dataset included only interactions between immune cells and relevant cytokines.

We classified interactions into:

- Direct cell-cell interactions, where both the source and target were immune cells.
- One-step mediated interactions, where one cell secretes a cytokine and another cell responds to that cytokine.

To assign each inferred cell-cell interaction a net sign, we applied the following rule:

- If either step involved Inhibit or Kill, the resulting interaction was marked as Negative.
- Otherwise, if both steps involved Activate, Survive, or Secrete, the interaction was marked as Positive.

We also removed ubiquitous cytokine mediators such as IFN-γ and TGF-β, which otherwise dominate the network structure.

#### 6.2 Graph Flattening and Annotating

We then constructed a flattened graph, where each cell-cell edge is represented once, and its regulatory nature is captured by two binary weights:

- A positive weight is assigned if there exists at least one path (direct or mediated) between the source and target with an Activate or Survive action.
- A negative weight is assigned if there exists at least one path with a Kill or Inhibit action.

This was implemented using a function that aggregated all interactions between each source-target cell pair. For each pair:

- The corresponding subset of the dataset was extracted.
- Boolean masks were applied to identify activating or inhibitory actions.
- A positive weight was set to 1 if any positive interaction was present, and a negative weight was set to 1 if any inhibitory interaction was present.
- The full list of contributing interactions was stored for downstream inspection.

This flattened, annotated, and weighted graph provided a biologically grounded and technically consistent platform for identifying structural motifs underlying immune cell dynamics.

#### 6.3 Motif Identification and Enrichment Analysis

To determine whether the excitable cell-cell motif identified in Section 5 is present in the immune network, we performed a subgraph search for the characteristic architecture consisting of:

- A self-activating effector cell X (positive self-loop),
- A positive interaction from X to an inhibitor cell Y,
- A negative interaction from Y back to X.

We implemented this search using the Labeled Adjacency Descriptor (LAD) subgraph matching algorithm from igraph. We defined a two-node directed motif with three edges: a self-loop on node A (X → X), a forward edge (X → Y), and a feedback edge (Y → X). All edges were required to match specific action patterns:

- The self-loop and X→Y edge had to include a **positive weight** (indicating activation or survival),
- The Y→X feedback edge had to include a **negative weight** (indicating inhibition or killing).

For each motif match, we confirmed that the weights were consistent with the required pattern. We also preserved the original edge annotations (including intermediary cytokines and immune processes) to allow biological interpretation of the matched circuits.

#### 6.4 Statistical Enrichment of Excitable Motifs in the Cell-Cell Graph

To determine whether the excitable cell-cell circuit motif was overrepresented in the immune interaction network compared to random expectation, we generated a null distribution of motif counts using randomized versions of the cell-cell network. We constructed these randomized graphs using a custom edge-rewiring algorithm that preserves key biological and structural properties of the original graph while disrupting specific motif arrangements.

The rewiring algorithm began with the cell-cell interaction graph described in section 6.1 above. Each edge was labeled as Positive or Negative depending on whether it included at least one activating (Activate or Survive) or inhibitory (Inhibit or Kill) interaction, respectively. These interaction signs were preserved during the rewiring process.

In each randomization:

1. We sampled two edges at random from the graph and proposed swapping their targets to generate new directed edges.
2. The proposed swap was only accepted if it did not introduce duplicate edges of the same type. This process was repeated for a number of swaps proportional to the total number of edges in the network (by default, one swap per edge).
3. After all swaps, we reconstructed a flattened version of the randomized network by aggregating all rewired edges as described in section 6.2

This procedure yielded randomized graphs that:

- Preserved the total number of edges,
- Preserved the number of positive and negative interactions,
- Maintained the degree distribution across nodes.

We generated 1000 such randomized networks and performed the same LAD-based subgraph search on each to count the number of excitable motifs. The distribution of motif counts across randomizations served as a null model against which the observed motif count (21) in the actual immune network was compared. The excitable motif was found to be significantly enriched (p = 0.04), suggesting that this specific regulatory architecture is not a statistical artifact, but rather a recurrent design principle in immune cell population dynamics.

### 7. Modeling Pro-Inflammatory Markers and IL-10 Dynamics in COVID-19 Patients

#### 7.1 Data Processing and Feature Construction

We analyzed longitudinal plasma cytokine measurements from 18 SARS–CoV–2–infected patients, recorded daily after inoculation. Data was imported from ref ^39^ and filtered to include only time points with days post-infection (dpi > 0). To characterize pro-inflammatory activity, we selected seven cytokines measured in the experiment, that were found in the motif search to participate in an excitable system with IL-10 as an inhibitor: IFN-α2a, IL-1α, IL-1β, TNF-α, IL-6, CCL4, and CCL2. For each patient and cytokine, values were min-normalized and converted to rank-normalized scores to allow aggregation across scales.

At each time point, we computed a rank-sum score across these seven cytokines to create a composite measure of the pro-inflammatory signal. IL-10 values were used directly to represent the anti-inflammatory response. To improve temporal resolution around dynamic peaks, we applied linear interpolation around the local maxima of both signals, generating two additional time points between each observed pair in a window of ±1 day surrounding the peaks. This produced a smoothed and temporally refined trajectory for each patient, restricted to the first 12 days post-infection. For patient number 10, the data point at day 7 was omitted due to a large deviation.

#### 7.2 Dynamical Model Structure

We modeled the interaction between pro-inflammatory signals and IL-10 using a two-dimensional system of ordinary differential equations (ODEs). The variable x(t) denotes the composite pro-inflammatory rank-sum score, while y(t) represents IL-10 concentration. The dynamics of these two variables were fitted to the following model:

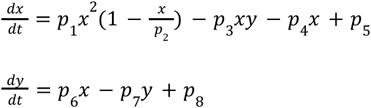

We implemented this system numerically using the RK45 solver from scipy.integrate.solve_ivp, with initial conditions derived from the first data point for each patient. To prevent numerical instabilities, simulated values below zero were truncated.

#### 7.3 Parameter Estimation via Weighted Least Squares

For each patient, we fitted the model parameters by minimizing the squared residuals between the simulated trajectories and the observed cytokine time series. The optimization was performed using the scipy.optimize.least_squares algorithm with bounds enforcing non-negativity. A weight matrix was applied to emphasize the peak pro-inflammatory and IL-10 values, which are most critical for determining pulse timing and amplitude.

#### 7.4 Bayesian Inference and Uncertainty Quantification

To quantify uncertainty and assess the robustness of the model fits, we used Markov Chain Monte Carlo (MCMC) to sample the posterior distribution of model parameters. We defined a log-likelihood assuming Gaussian measurement noise with standard deviation σ = 0.05 and added peak-emphasized weights to the squared error term. A uniform prior between 0 and 50 was placed on all parameters.

We ran the MCMC using the emcee ensemble sampler^68^ with 24 walkers and 1000 steps, initialized around the least-squares solution. The first 200 steps were discarded as burn-in, and the chains were thinned by a factor of 10 to reduce autocorrelation. From the resulting posterior samples, we computed the mean and 90% credible intervals for each parameter.

#### 7.5 Posterior Predictive Simulation and Model Validation

To evaluate the predictive accuracy of the model while accounting for uncertainty in parameter estimates, we performed posterior predictive simulations. For each patient, we selected 100 parameter samples from the posterior distribution obtained by MCMC and simulated the corresponding cytokine trajectories over time. We then computed the 2.5th, 50th (median), and 97.5th percentiles across these trajectories at each time point to construct 95% credible intervals for both the pro-inflammatory signal and IL-10 levels.

Model performance was assessed by comparing the observed patient trajectories to the predicted median and credible intervals. We computed the posterior predictive coverage, defined as the fraction of observed data points falling within the 95% credible intervals.

To quantitatively assess model fit quality, we calculated both the standard R^2^ and a weighted R^2^ score. The weighted R2 was designed to emphasize the accurate reproduction of key features in the dynamics, particularly the timing and amplitude of pro-inflammatory markers and IL-10 peaks, which reflect the system’s excitable behavior. For each patient, we defined a weight matrix where time points corresponding to the peak of the pro-inflammatory markers response and the IL-10 response were assigned higher weights (value 10), while all other points were assigned a weight 1.

The weighted R^2^ was computed as:

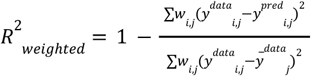

where *w* _*i,j*_ is the weight at timepoint i for variable j, and 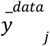 is the mean value of the observed data for variable j.

In most patients, the weighted R^2^ exceeded the unweighted value, and in the majority of cases was above 0.8, indicating that the excitable circuit model captured both the qualitative shape and quantitative features of the immune response. The full list of model validation results can be found in supplementary table 2.

### 8. Circuit to target and sensitivity analysis

#### 8.1 Model equations and parameters

To simplify the analysis, we worked with a reduced version of the basic architecture of the excitable cell-cell circuit found in the screening:

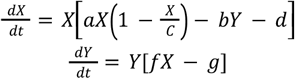

The parameters have the following units: 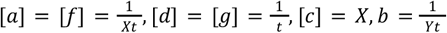.

#### 8.2 Threshold analysis

The threshold was calculated as the minimal X in which 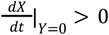, which is given by:

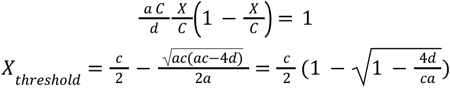

Thus, the threshold depends only on the parameters a,c, and d. Sensitivity analysis was performed by setting a set of parameters as described in Fig.ure 6 and varying each of the parameters individually. The threshold in the above equation was numerically calculated for each of the parameter values. Then a first-order estimate of the derivative of the log was taken by finite differences of order 1. The parameter range for the analysis can be found in the SI (Supplementary table 4).

#### 8.3 Pulse amplitude analysis

The amplitude was calculated by integrating the ODEs forward in time, with initial conditions given by the 110% threshold calculated in part 7.2 for each set of parameters. Each integration was carried out until t=1 or until the maximum level of X was reached. The amplitude was taken to be the maximum level of X reached. Integration in time was done using the Euler method with dt=0.001. Sensitivity to each of the parameters was taken as described in 7.2. The parameter range for the analysis can be found in the SI.

#### 8.4 Refractory period analysis

The ODEs were integrated forward in time, with initial conditions of *X* = 10, *Y* = 0. 1. To calculate the refractory period, we first retrieved the maximum of Y for the integration. We then calculated the first time (after the peak) in which Y levels decreased below 1% of the maximum. The integration was carried out until t=10 or until Y levels decreased below 1% of the maximum. Integration in time was done using the Euler method with dt=0.001. Sensitivity to each of the parameters was taken as described in 7.2. The parameter range for the analysis can be found in the SI.

#### 8.5 Phase Diagram

The model presented in 8.1 was nondimensionalized to the form:

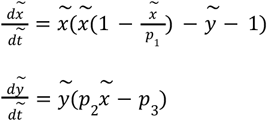

With:

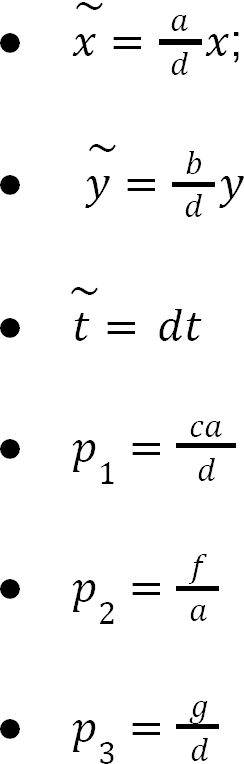

With *p*_3_ = 1 as we assume no timescale separation in this problem. Excitability was defined as the range of parameters in which the nullcline 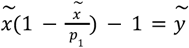 crosses the x axis in two points, and that there is no other stable fixed point other than the origin: this regime was in the regime 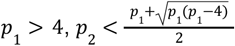. In the regime *p* _1_ < 4, parabole nullcline doesnt cross the x axis, resulting in only the origin being a stable fixed point; in the regime 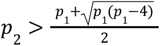 a new stable fixed point is created at 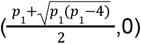.

---

**Supplementary table 1.**
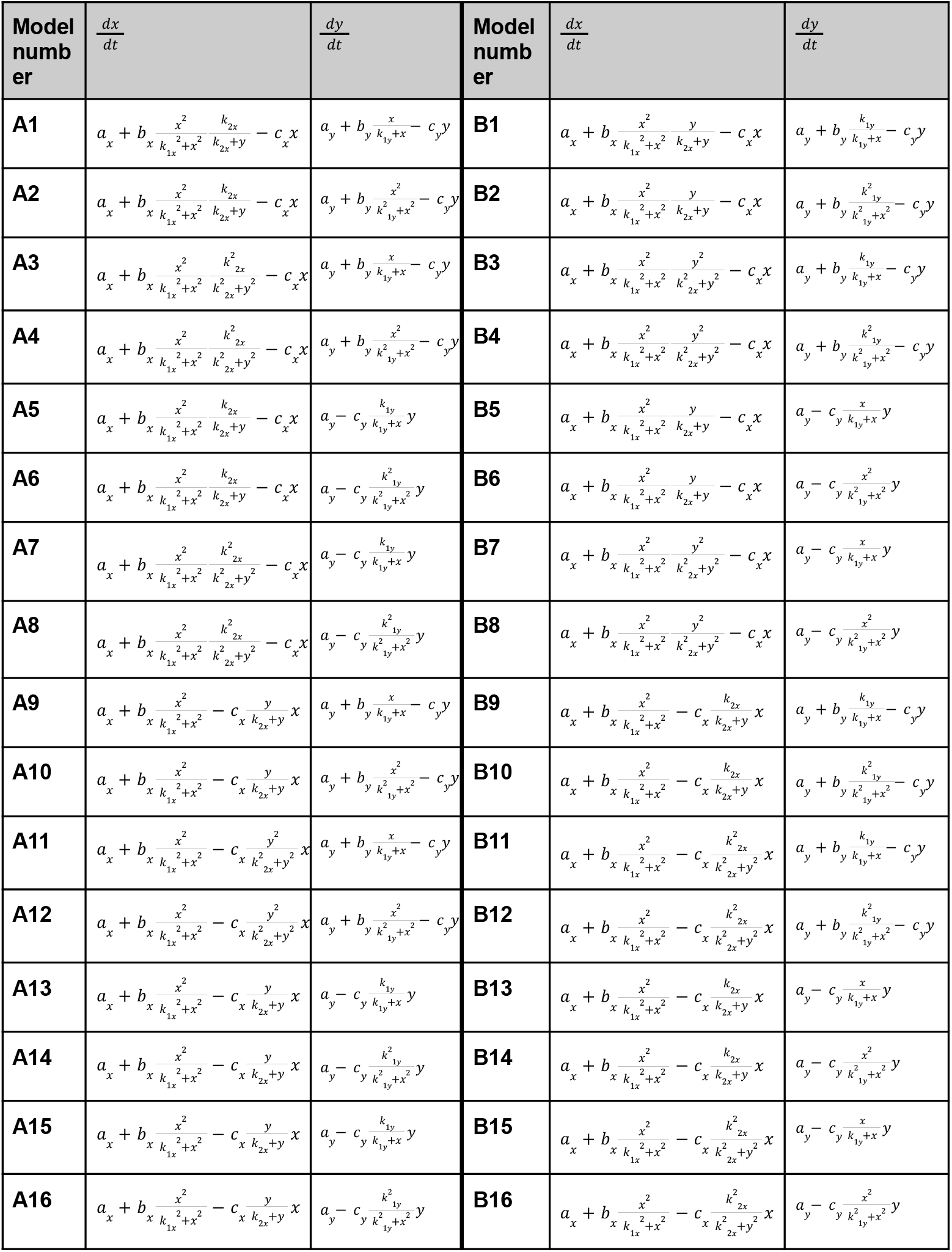
Circuit equations for the 32 excitable cytokine circuits.

**Supplementary table 2.**
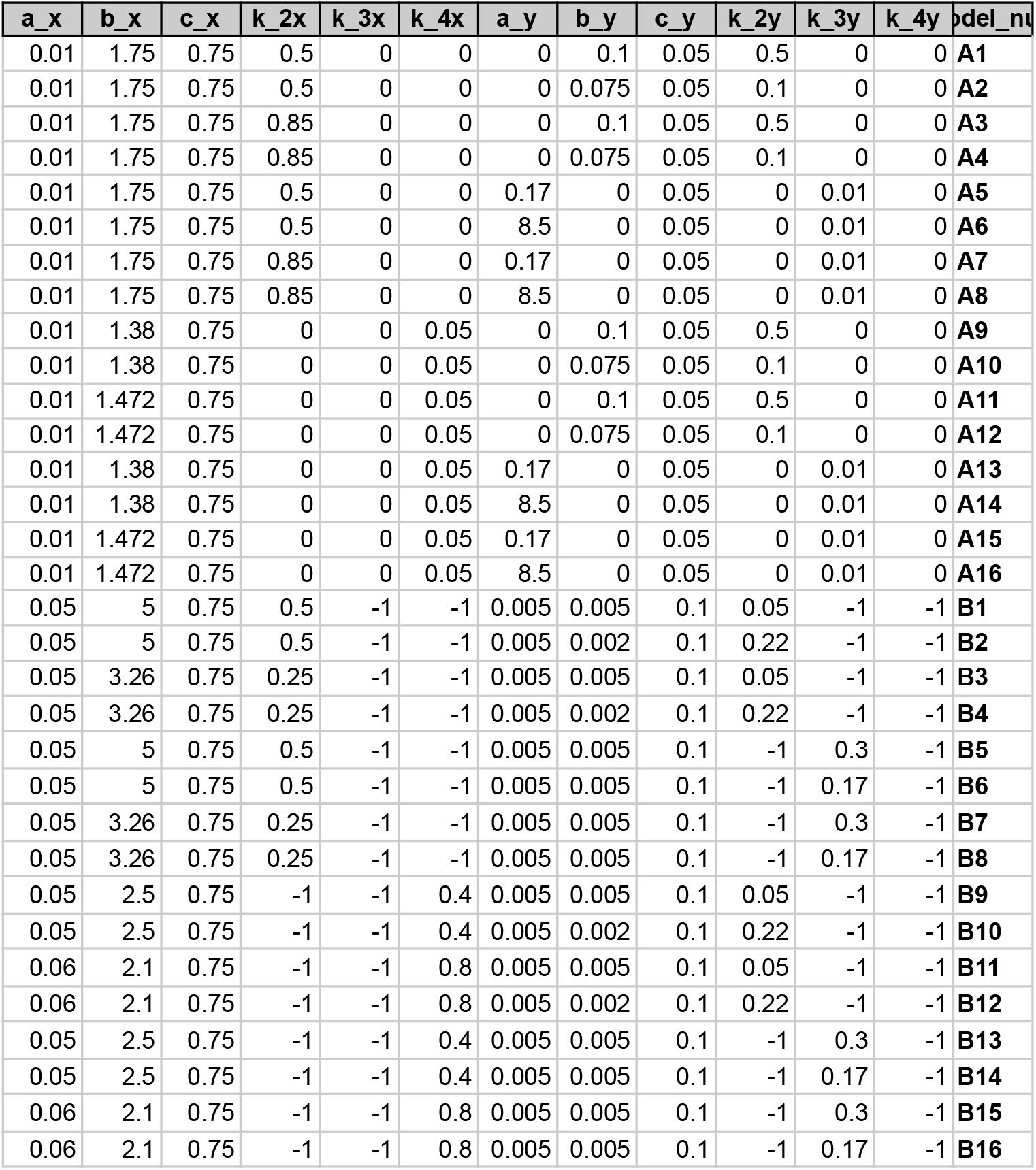
Parameters used to compare models.

**Supplementary table 3.**
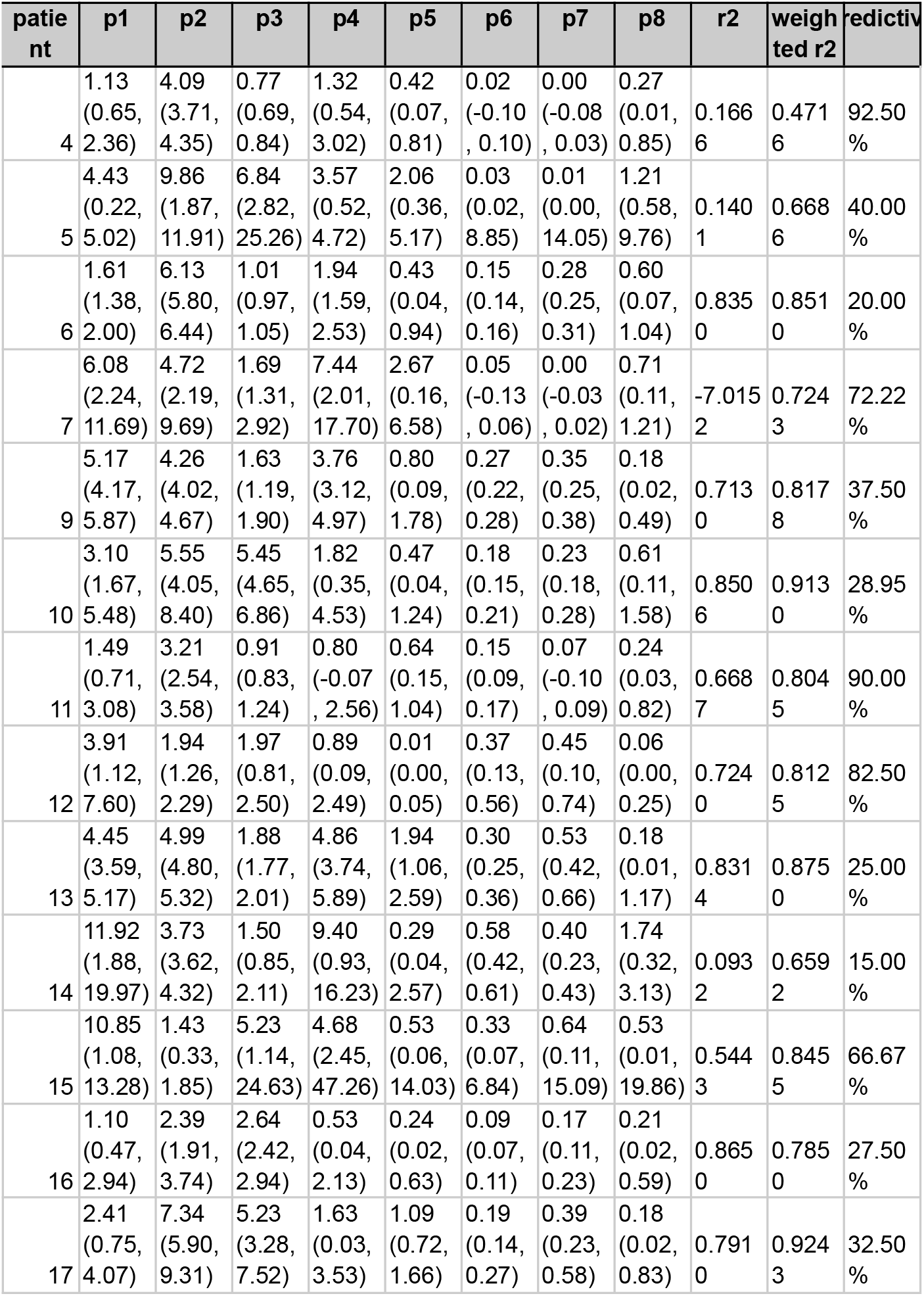
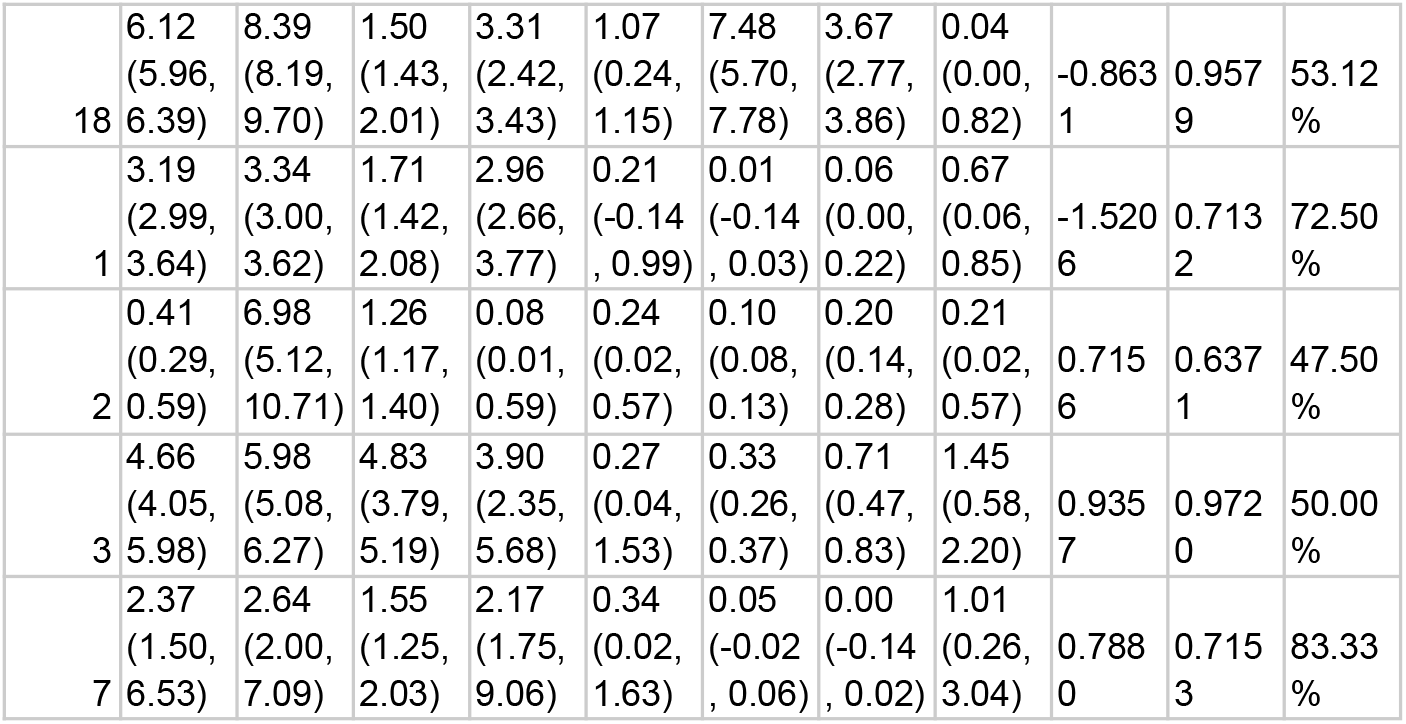
Best fit parameters for pro inflammatory-IL10 circuit.

**Supplementary table 4.**
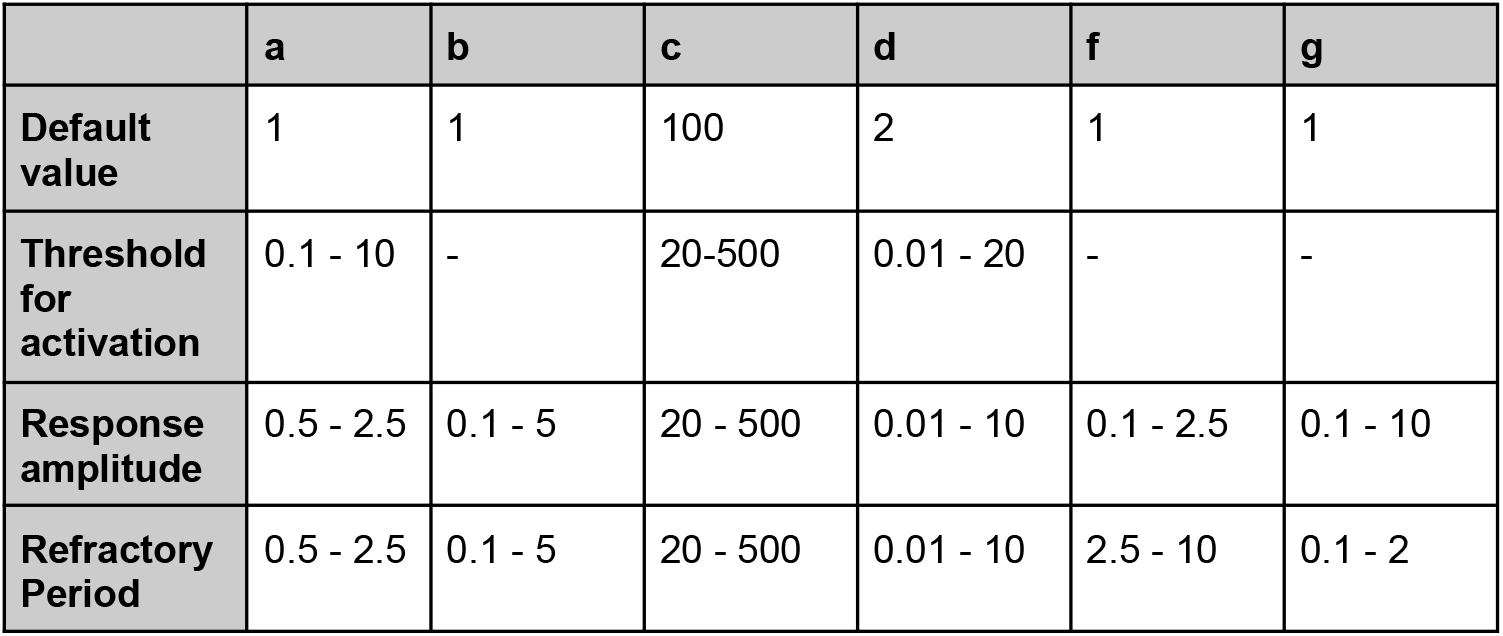
Parameter range for sensitivity analysis.

**Supplementary Figure 1.**
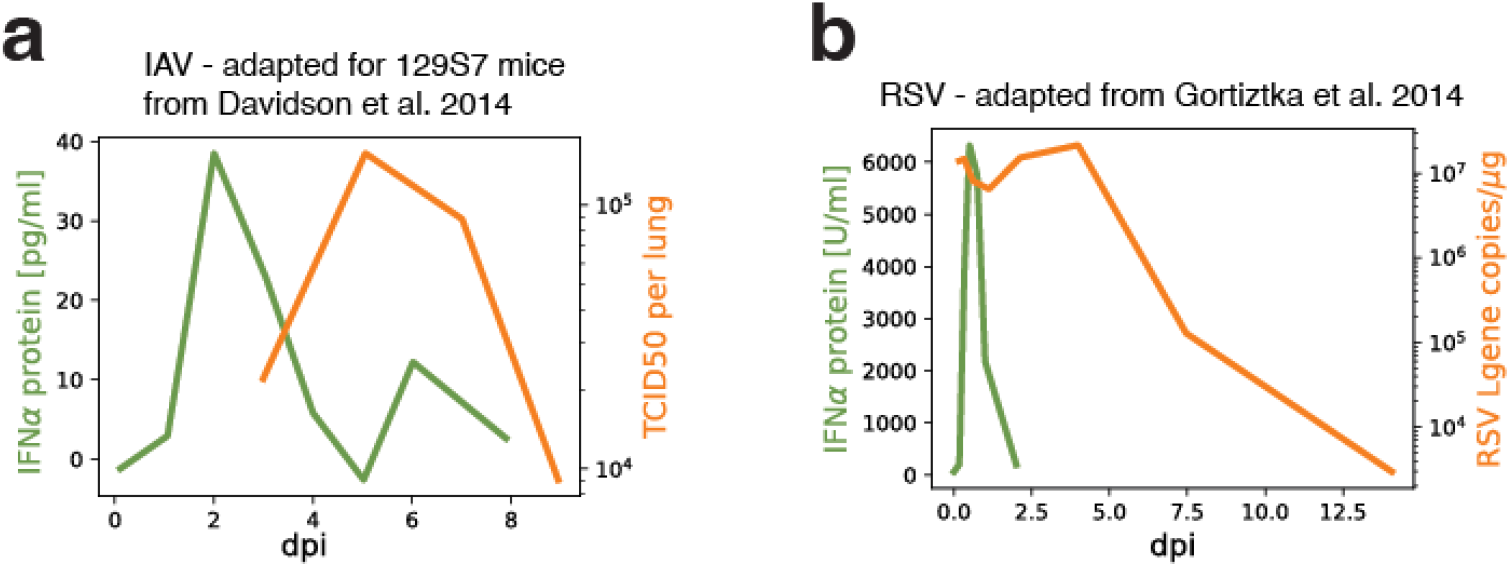
Type I interferon pulse ends before viral load decreases in mice infected with IAV and RSV. a) Type I interferon levels (green) vs. viral load (orange) measured in 129S7 mice infected with IAV over the first 8 days post-infection (dpi). Adapted from Davidson et al. 2014. b) Type I interferon levels (green) vs. viral load (orange) measured in mice infected with RSV over the first 14 days post-infection. Adapted from Goritzka et al. 2014.

**Supplementary figure 2.**
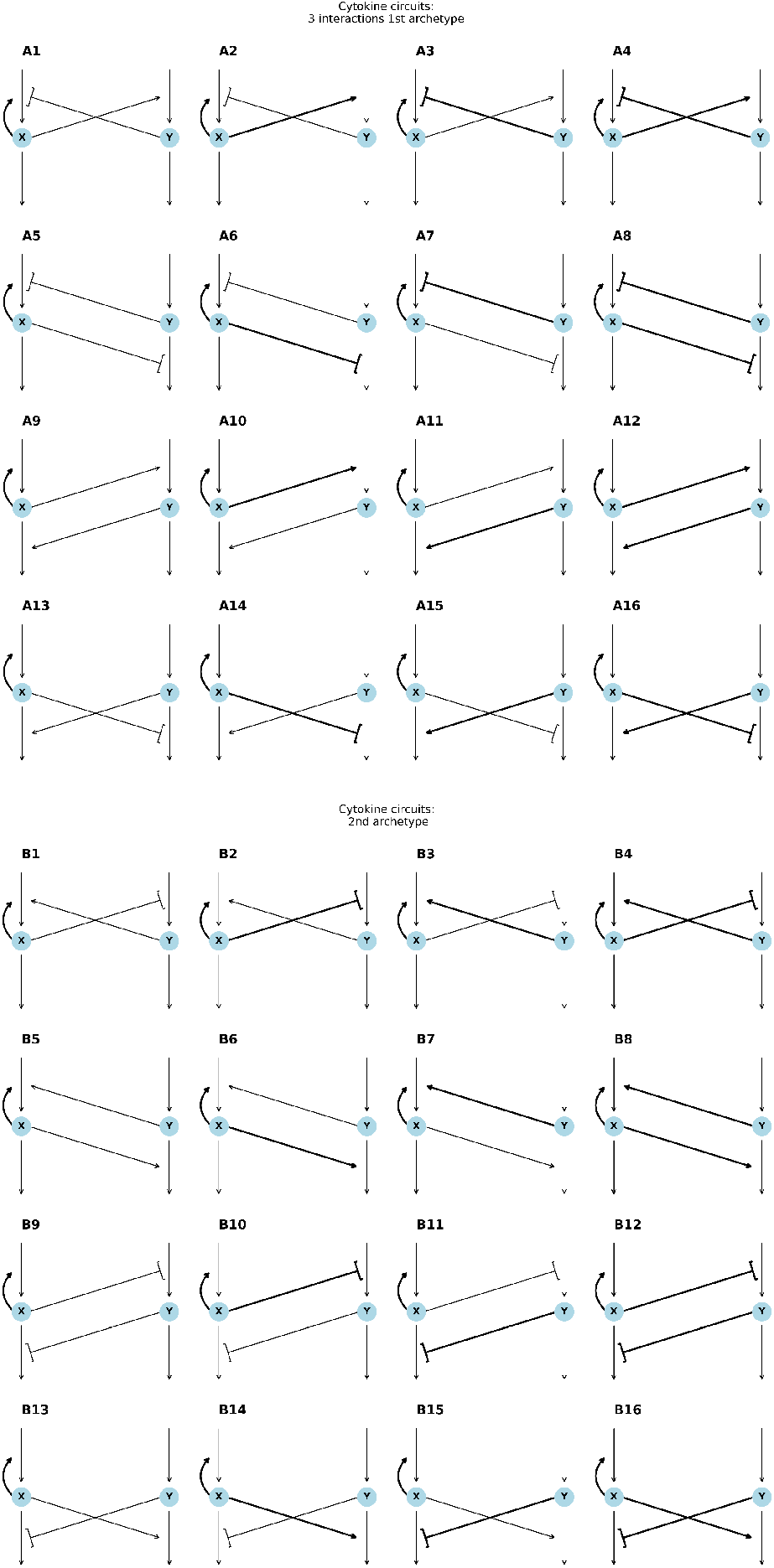
All excitable cytokine circuit topologies. This list is corresponding with supplementary table 1.

**Supplementary figure 3.**
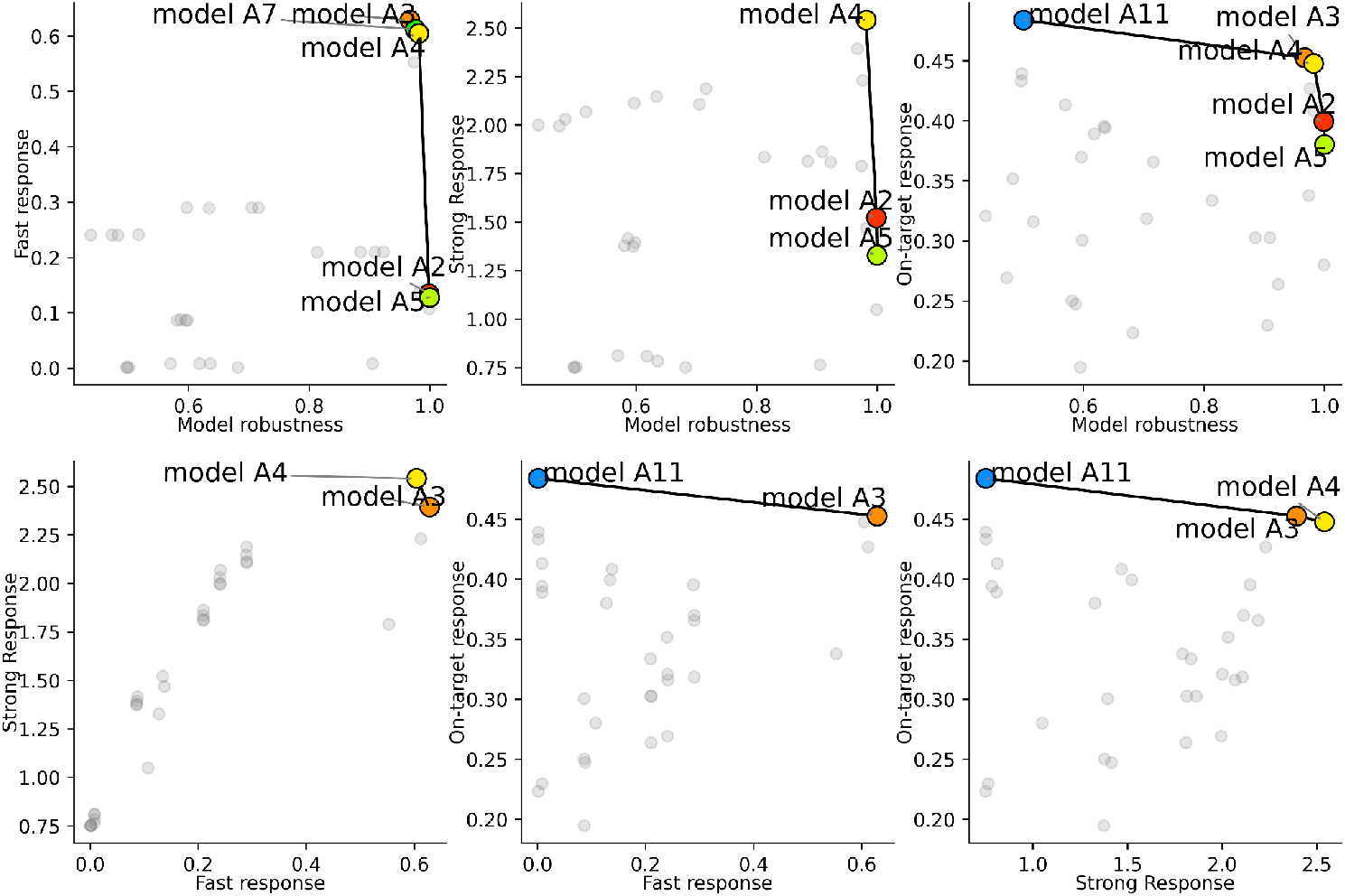
Pareto fronts of models in all metrics.

**Supplementary Figure 4.**
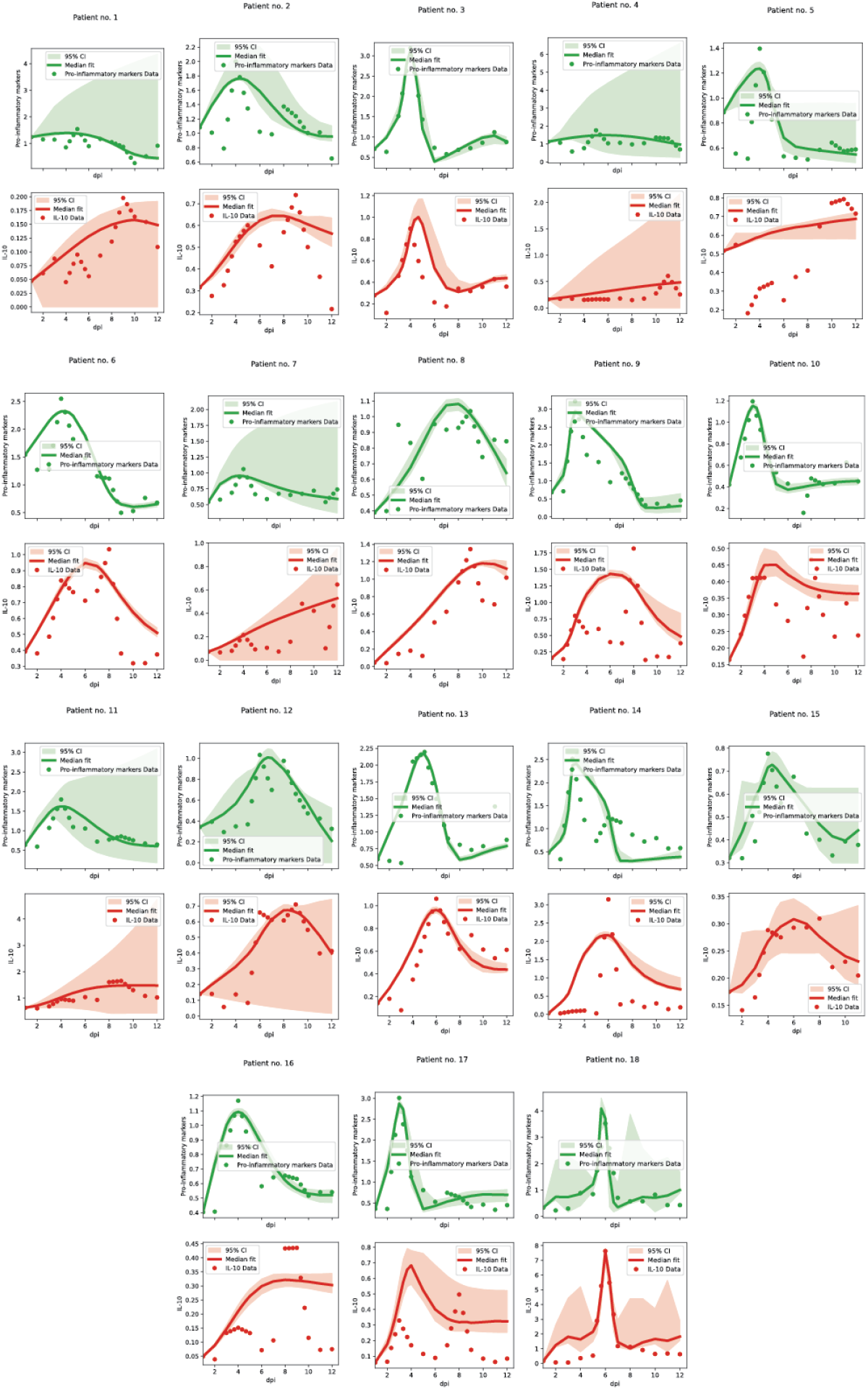
Pro inflammatory cytokines-IL10 model fit to all patients.

**Supplementary Figure 5.**
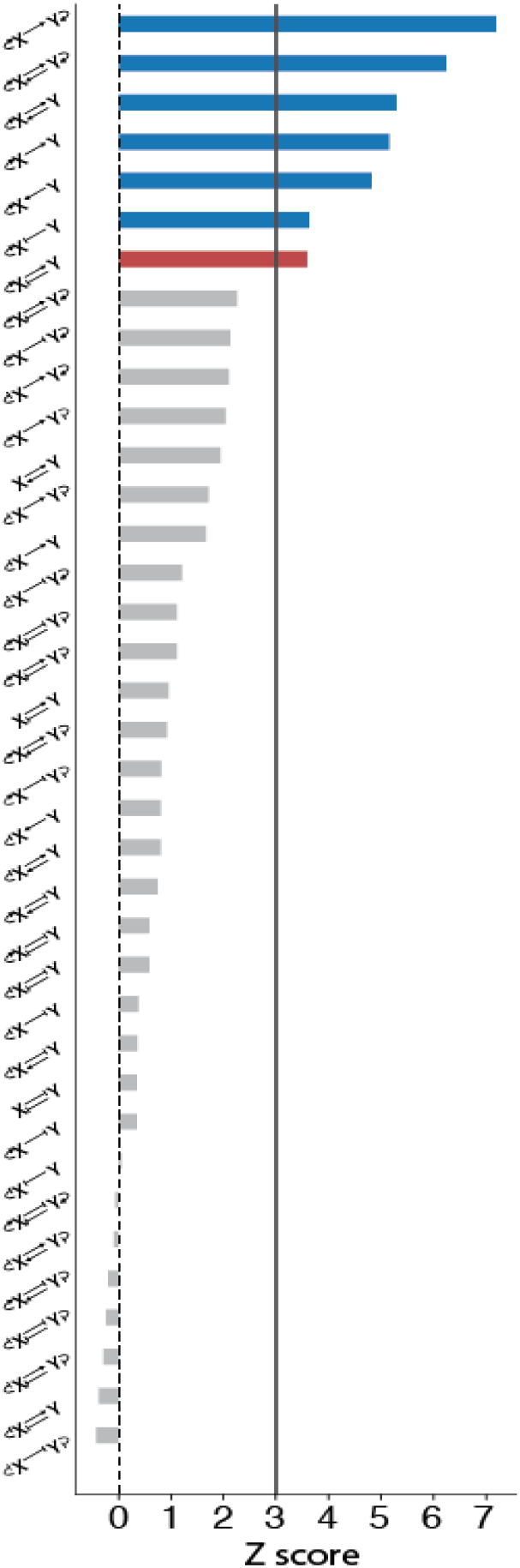
Motifs distribution in the cytokine network. X-axis is the z score of each motif. Blue bars are significantly enriched (z-score>3). The red bar is the excitable motif.

## Notes

### Competing Interest Statement

The authors have declared no competing interest.

### Funding

- Azrieli Foundation

## References

1. Alon, U. An Introduction to Systems Biology: Design Principles of Biological Circuits. (Chapman and Hall/CRC, Second edition. | Boca Raton, Fla. : CRC Press, [2019], 2019). doi:10.1201/9780429283321.

2. Perelson, A. S. & Weisbuch, G. Immunology for physicists. Rev. Mod. Phys. 69, 1219–1268 (1997).

3. Chakraborty, A. K. A Perspective on the Role of Computational Models in Immunology. Annu. Rev. Immunol. 35, 403–439 (2017).

4. Altan-Bonnet, G., Mora, T. & Walczak, A. M. Quantitative immunology for physicists. Phys. Rep. 849, 1–83 (2020).

5. Hodgkin, P. D., Dowling, M. R. & Duffy, K. R. Why the immune system takes its chances with randomness. Nat. Rev. Immunol. 14, 711–711 (2014).

6. Davis, M. M., Tato, C. M. & Furman, D. Systems immunology: just getting started. Nat. Immunol. 18, 725–732 (2017).

7. Jerne, N. K. Towards a network theory of the immune system. Ann. Immunol. 125C, 373–389 (1974).

8. Perelson, A. S. Immune Network Theory. Immunol. Rev. 110, 5–36 (1989).

9. Medzhitov, R. & Janeway, C. Innate immunity. N. Engl. J. Med. 343, 338–344 (2000).

10. Hoffmann, A., Levchenko, A., Scott, M. L. & Baltimore, D. The IκB-NF-κB Signaling Module: Temporal Control and Selective Gene Activation. Science 298, 1241–1245 (2002).

11. Ashkenazi, A. & Dixit, V. M. Death Receptors: Signaling and Modulation. Science 281, 1305–1308 (1998).

12. Nowak, M. A. & Bangham, C. R. M. Population Dynamics of Immune Responses to Persistent Viruses. Science 272, 74–79 (1996).

13. Frank, S. A. Immune Response to Parasitic Attack: Evolution of a Pulsed Character. J. Theor. Biol. 219, 281–290 (2002).

14. Rand, U. et al. Multi-layered stochasticity and paracrine signal propagation shape the type-I interferon response. Mol. Syst. Biol. 8, 584 (2012).

15. Medzhitov, R. Origin and physiological roles of inflammation. Nature 454, 428–435 (2008).

16. Kaech, S. M. & Wherry, E. J. Heterogeneity and cell-fate decisions in effector and memory CD8+ T cell differentiation during viral infection. Immunity 27, 393–405 (2007).

17. Altan-Bonnet, G. & Germain, R. N. Modeling T Cell Antigen Discrimination Based on Feedback Control of Digital ERK Responses. PLOS Biol. 3, e356 (2005).

18. Lebel, Y., Milo, T., Bar, A., Mayo, A. & Alon, U. Excitable dynamics of flares and relapses in autoimmune diseases. iScience 26, 108084 (2023).

19. Osojnik, A., Gaffney, E. A., Davies, M., Yates, J. W. T. & Byrne, H. M. Identifying and characterising the impact of excitability in a mathematical model of tumour-immune interactions. J. Theor. Biol. 501, 110250 (2020).

20. Somer, J., Mannor, S. & Alon, U. Temporal tissue dynamics from a single snapshot. 2024.04.22.590503 Preprint at 10.1101/2024.04.22.590503 (2024).

21. Weiss, C. O. & Vilaseca, R. Dynamics of lasers. NASA STIRecon Tech. Rep. A 92, 39875 (1991).

22. Meron, E. Pattern formation in excitable media. Phys. Rep. 218, 1–66 (1992).

23. Krauskopf, B., Schneider, K., Sieber, J., Wieczorek, S. & Wolfrum, M. Excitability and self-pulsations near homoclinic bifurcations in semiconductor laser systems. Opt. Commun. 215, 367–379 (2003).

24. Lindner, B., García-Ojalvo, J., Neiman, A. & Schimansky-Geier, L. Effects of noise in excitable systems. Phys. Rep. 392, 321–424 (2004).

25. Hodgkin, A. L. & Huxley, A. F. A quantitative description of membrane current and its application to conduction and excitation in nerve. J. Physiol. 117, 500–544 (1952).

26. FitzHugh, R. Mathematical models of threshold phenomena in the nerve membrane. Bull. Math. Biophys. 17, 257–278 (1955).

27. Nagumo, J., Arimoto, S. & Yoshizawa, S. An Active Pulse Transmission Line Simulating Nerve Axon. Proc. IRE 50, 2061–2070 (1962).

28. Keener, J. & Sneyd, J. Mathematical Physiology. vol. 8 (Springer New York, New York, NY, 1998).

29. Gerstner, W. & Kistler, W. M. Spiking Neuron Models: Single Neurons, Populations, Plasticity. (Cambridge University Press, Cambridge, 2002). doi:10.1017/CBO9780511815706.

30. Izhikevich, E. M. Neural excitability, spiking and bursting. Int. J. Bifurc. Chaos Appl. Sci. Eng. 10, 1171–1266 (2000).

31. Othmer, Hans. G. The Dynamics of Forced Excitable Systems. in Nonlinear Wave Processes in Excitable Media (eds. Holden, A. V., Markus, M. & Othmer, H.G.) 213–231 (Springer US, Boston, MA, 1991). doi:10.1007/978-1-4899-3683-7_21.

32. Strogatz, S. H. Nonlinear Dynamics and Chaos. (CRC Press, 2018). doi:10.1201/9780429492563.

33. Gutkin, B. S. & Ermentrout, G. B. Dynamics of Membrane Excitability Determine Interspike Interval Variability: A Link Between Spike Generation Mechanisms and Cortical Spike Train Statistics. Neural Comput. 10, 1047–1065 (1998).

34. Savageau, M. A. The Behavior of Intact Biochemical Control Systems*. in Current Topics in Cellular Regulation (eds. Horecker, B.L. & Stadtman, E.R.) vol. 6 63–130 (Academic Press, 1972).

35. Savageau, M. A. Comparison of classical and autogenous systems of regulation in inducible operons. Nature 252, 546–549 (1974).

36. Alves, R. & Savageau, M. A. Extending the method of mathematically controlled comparison to include numerical comparisons. Bioinformatics 16, 786–798 (2000).

37. Adler, M., Szekely, P., Mayo, A. & Alon, U. Optimal Regulatory Circuit Topologies for Fold-Change Detection. Cell Syst. 4, 171–181.e8 (2017).

38. Atallah, M. B. et al. ImmunoGlobe: enabling systems immunology with a manually curated intercellular immune interaction network. BMC Bioinformatics 21, 346 (2020).

39. Wagstaffe, H. R. et al. Mucosal and systemic immune correlates of viral control after SARS-CoV-2 infection challenge in seronegative adults. Sci. Immunol. 9, eadj9285 (2024).

40. Tan, J., Pan, R., Qiao, L., Zou, X. & Pan, Z. Modeling and Dynamical Analysis of Virus-Triggered Innate Immune Signaling Pathways. PLoS ONE 7, e48114 (2012).

41. Davidson, S., Crotta, S., McCabe, T. M. & Wack, A. Pathogenic potential of interferon αβ in acute influenza infection. Nat. Commun. 5, 3864 (2014).

42. Goritzka, M. et al. Alpha/Beta Interferon Receptor Signaling Amplifies Early Proinflammatory Cytokine Production in the Lung during Respiratory Syncytial Virus Infection. J. Virol. 88, 6128–6136 (2014).

43. Heim, M. H. Innate immunity and HCV. J. Hepatol. 58, 564–574 (2013).

44. Shin, E.-C., Sung, P. S. & Park, S.-H. Immune responses and immunopathology in acute and chronic viral hepatitis. Nat. Rev. Immunol. 16, 509–523 (2016).

45. Zhou, X. et al. Circuit Design Features of a Stable Two-Cell System. Cell 172, 744–757.e17 (2018).

46. Mueller, S. N. & Rouse, B. T. Immune responses to viruses. Clin. Immunol. 421–431 (2008) doi:10.1016/B978-0-323-04404-2.10027-2.

47. Rouse, B. T., Sarangi, P. P. & Suvas, S. Regulatory T cells in virus infections. Immunol. Rev. 212, 272–286 (2006).

48. Velez de Mendizabal, N. et al. Modeling the effector - Regulatory T cell cross-regulation reveals the intrinsic character of relapses in Multiple Sclerosis. BMC Syst. Biol. 5, 1–15 (2011).

49. Butler, T. C., Kardar, M. & Chakraborty, A. K. Quorum sensing allows T cells to discriminate between self and nonself. Proc. Natl. Acad. Sci. U. S. A. 110, 11833–11838 (2013).

50. Sojka, D. K., Huang, Y. H. & Fowell, D. J. Mechanisms of regulatory T-cell suppression - A diverse arsenal for a moving target. Immunology 124, 13–22 (2008).

51. Rosser, E. C. & Mauri, C. Regulatory B Cells: Origin, Phenotype, and Function. Immunity 42, 607–612 (2015).

52. Michaud, D., Steward, C. R., Mirlekar, B. & Pylayeva-Gupta, Y. Regulatory B cells in cancer. Immunol. Rev. 299, 74–92 (2021).

53. Zi, Z. Sensitivity analysis approaches applied to systems biology models. IET Syst. Biol. 5, 336–336 (2011).

54. Blanco-Melo, D. et al. Imbalanced Host Response to SARS-CoV-2 Drives Development of COVID-19. Cell 181, 1036–1045.e9 (2020).

55. Tagliabue, A., McCoy, J. L. & Herberman, R. B. Refractoriness to Migration Inhibitory Factor of Macrophages of LPS Nonresponder Mouse Strains. J. Immunol. 121, 1223–1226 (1978).

56. Fahmi, H. & Chaby, R. Selective refractoriness of macrophages to endotoxin-induced production of tumor necrosis factor, elicited by an autocrine mechanism. J. Leukoc. Biol. 53, 45–52 (1993).

57. Galoppin, L. et al. Nonspecific refractoriness to adenylyl cyclase stimulation in alveolar macrophages from infants with recurrent bronchiolitis. J. Allergy Clin. Immunol. 93, 885–890 (1994).

58. Zhang, Z., Penn, R., Barclay, W. S. & Giotis, E. S. Naïve Human Macrophages Are Refractory to SARS-CoV-2 Infection and Exhibit a Modest Inflammatory Response Early in Infection. Viruses 14, 441 (2022).

59. Duthoit, C. T., Nguyen, P. & Geiger, T. L. Antigen nonspecific suppression of T cell responses by activated stimulation-refractory CD4+ T cells. J. Immunol. Baltim. Md 1950 172, 2238–2246 (2004).

60. Appleman, L. J. & Boussiotis, V. A. T cell anergy and costimulation. Immunol. Rev. 192, 161–180 (2003).

61. Viola, A. & Lanzavecchia, A. T Cell Activation Determined by T Cell Receptor Number and Tunable Thresholds. Science 273, 104–106 (1996).

62. Halova, I. et al. Changing the threshold-Signals and mechanisms of mast cell priming. Immunol. Rev. 282, 73–86 (2018).

63. Gilfillan, A. M., Peavy, R. D. & Metcalfe, D. D. Amplification mechanisms for the enhancement of antigen-mediated mast cell activation. Immunol. Res. 43, 15–24 (2009).

64. Miyara, S. et al. Cold and hot fibrosis define clinically distinct cardiac pathologies. Cell Syst. 16, (2025).

65. Stearns, S. C. & Medzhitov, R. Evolutionary Medicine. (Oxford University Press, Oxford, 2024).

66. Zulko. Zulko/ddeint. (2024).

67. Virtanen, P. et al. SciPy 1.0: fundamental algorithms for scientific computing in Python. Nat. Methods 17, 261–272 (2020).

68. Foreman-Mackey, D., Hogg, D. W., Lang, D. & Goodman, J. emcee: The MCMC Hammer. Publ. Astron. Soc. Pac. 125, 306–312 (2013).

