## Supplementary material for "Excitability as a Design Principle in the Immune System"

**Supplementary table 1. Circuit equations for the 32 excitable cytokine circuits.**

| Model number | $\frac{dx}{dt}$ | $\frac{dy}{dt}$ | Model number | $\frac{dx}{dt}$ | $\frac{dy}{dt}$ |
|---|---|---|---|---|---|
| A1 | $a_x + b_x \frac{x^2}{k_{1x}^2+x^2}\frac{k_{2x}}{k_{2x}+y} - c_x x$ | $a_y + b_y \frac{x}{k_{1y}+x} - c_y y$ | B1 | $a_x + b_x \frac{x^2}{k_{1x}^2+x^2}\frac{y}{k_{2x}+y} - c_x x$ | $a_y + b_y \frac{k_{1y}}{k_{1y}+x} - c_y y$ |
| A2 | $a_x + b_x \frac{x^2}{k_{1x}^2+x^2}\frac{k_{2x}}{k_{2x}+y} - c_x x$ | $a_y + b_y \frac{x^2}{k_{1y}^2+x^2} - c_y y$ | B2 | $a_x + b_x \frac{x^2}{k_{1x}^2+x^2}\frac{y}{k_{2x}+y} - c_x x$ | $a_y + b_y \frac{k_{1y}^2}{k_{1y}^2+x^2} - c_y y$ |
| A3 | $a_x + b_x \frac{x^2}{k_{1x}^2+x^2}\frac{k_{2x}^2}{k_{2x}^2+y^2} - c_x x$ | $a_y + b_y \frac{x}{k_{1y}+x} - c_y y$ | B3 | $a_x + b_x \frac{x^2}{k_{1x}^2+x^2}\frac{y^2}{k_{2x}^2+y^2} - c_x x$ | $a_y + b_y \frac{k_{1y}}{k_{1y}+x} - c_y y$ |
| A4 | $a_x + b_x \frac{x^2}{k_{1x}^2+x^2}\frac{k_{2x}^2}{k_{2x}^2+y^2} - c_x x$ | $a_y + b_y \frac{x^2}{k_{1y}^2+x^2} - c_y y$ | B4 | $a_x + b_x \frac{x^2}{k_{1x}^2+x^2}\frac{y^2}{k_{2x}^2+y^2} - c_x x$ | $a_y + b_y \frac{k_{1y}^2}{k_{1y}^2+x^2} - c_y y$ |
| A5 | $a_x + b_x \frac{x^2}{k_{1x}^2+x^2}\frac{k_{2x}}{k_{2x}+y} - c_x x$ | $a_y - c_y \frac{k_{1y}}{k_{1y}+x} y$ | B5 | $a_x + b_x \frac{x^2}{k_{1x}^2+x^2}\frac{y}{k_{2x}+y} - c_x x$ | $a_y - c_y \frac{x}{k_{1y}+x} y$ |
| A6 | $a_x + b_x \frac{x^2}{k_{1x}^2+x^2}\frac{k_{2x}}{k_{2x}+y} - c_x x$ | $a_y - c_y \frac{k_{1y}^2}{k_{1y}^2+x^2} y$ | B6 | $a_x + b_x \frac{x^2}{k_{1x}^2+x^2}\frac{y}{k_{2x}+y} - c_x x$ | $a_y - c_y \frac{x^2}{k_{1y}^2+x^2} y$ |
| A7 | $a_x + b_x \frac{x^2}{k_{1x}^2+x^2}\frac{k_{2x}^2}{k_{2x}^2+y^2} - c_x x$ | $a_y - c_y \frac{k_{1y}}{k_{1y}+x} y$ | B7 | $a_x + b_x \frac{x^2}{k_{1x}^2+x^2}\frac{y^2}{k_{2x}^2+y^2} - c_x x$ | $a_y - c_y \frac{x}{k_{1y}+x} y$ |
| A8 | $a_x + b_x \frac{x^2}{k_{1x}^2+x^2}\frac{k_{2x}^2}{k_{2x}^2+y^2} - c_x x$ | $a_y - c_y \frac{k_{1y}^2}{k_{1y}^2+x^2} y$ | B8 | $a_x + b_x \frac{x^2}{k_{1x}^2+x^2}\frac{y^2}{k_{2x}^2+y^2} - c_x x$ | $a_y - c_y \frac{x^2}{k_{1y}^2+x^2} y$ |
| A9 | $a_x + b_x \frac{x^2}{k_{1x}^2+x^2} - c_x \frac{y}{k_{2x}+y} x$ | $a_y + b_y \frac{x}{k_{1y}+x} - c_y y$ | B9 | $a_x + b_x \frac{x^2}{k_{1x}^2+x^2} - c_x \frac{k_{2x}}{k_{2x}+y} x$ | $a_y + b_y \frac{k_{1y}}{k_{1y}+x} - c_y y$ |
| A10 | $a_x + b_x \frac{x^2}{k_{1x}^2+x^2} - c_x \frac{y}{k_{2x}+y} x$ | $a_y + b_y \frac{x^2}{k_{1y}^2+x^2} - c_y y$ | B10 | $a_x + b_x \frac{x^2}{k_{1x}^2+x^2} - c_x \frac{k_{2x}}{k_{2x}+y} x$ | $a_y + b_y \frac{k_{1y}^2}{k_{1y}^2+x^2} - c_y y$ |
| A11 | $a_x + b_x \frac{x^2}{k_{1x}^2+x^2} - c_x \frac{y^2}{k_{2x}^2+y^2} x$ | $a_y + b_y \frac{x}{k_{1y}+x} - c_y y$ | B11 | $a_x + b_x \frac{x^2}{k_{1x}^2+x^2} - c_x \frac{k_{2x}^2}{k_{2x}^2+y^2} x$ | $a_y + b_y \frac{k_{1y}}{k_{1y}+x} - c_y y$ |
| A12 | $a_x + b_x \frac{x^2}{k_{1x}^2+x^2} - c_x \frac{y^2}{k_{2x}^2+y^2} x$ | $a_y + b_y \frac{x^2}{k_{1y}^2+x^2} - c_y y$ | B12 | $a_x + b_x \frac{x^2}{k_{1x}^2+x^2} - c_x \frac{k_{2x}^2}{k_{2x}^2+y^2} x$ | $a_y + b_y \frac{k_{1y}^2}{k_{1y}^2+x^2} - c_y y$ |
| A13 | $a_x + b_x \frac{x^2}{k_{1x}^2+x^2} - c_x \frac{y}{k_{2x}+y} x$ | $a_y - c_y \frac{k_{1y}}{k_{1y}+x} y$ | B13 | $a_x + b_x \frac{x^2}{k_{1x}^2+x^2} - c_x \frac{k_{2x}}{k_{2x}+y} x$ | $a_y - c_y \frac{x}{k_{1y}+x} y$ |
| A14 | $a_x + b_x \frac{x^2}{k_{1x}^2+x^2} - c_x \frac{y}{k_{2x}+y} x$ | $a_y - c_y \frac{k_{1y}^2}{k_{1y}^2+x^2} y$ | B14 | $a_x + b_x \frac{x^2}{k_{1x}^2+x^2} - c_x \frac{k_{2x}}{k_{2x}+y} x$ | $a_y - c_y \frac{x^2}{k_{1y}^2+x^2} y$ |
| A15 | $a_x + b_x \frac{x^2}{k_{1x}^2+x^2} - c_x \frac{y}{k_{2x}+y} x$ | $a_y - c_y \frac{k_{1y}}{k_{1y}+x} y$ | B15 | $a_x + b_x \frac{x^2}{k_{1x}^2+x^2} - c_x \frac{k_{2x}^2}{k_{2x}^2+y^2} x$ | $a_y - c_y \frac{x}{k_{1y}+x} y$ |

| | | | | | |
|---|---|---|---|---|---|
| **A16** | $a_x + b_x \frac{x^2}{k_{1x}^2+x^2} - c_x \frac{y}{k_{2x}+y} x$ | $a_y - c_y \frac{k_{1y}^2}{k_{1y}^2+x^2} y$ | **B16** | $a_x + b_x \frac{x^2}{k_{1x}^2+x^2} - c_x \frac{k_{2x}^2}{k_{2x}^2+y^2} x$ | $a_y - c_y \frac{x^2}{k_{1y}^2+x^2} y$ |

**Supplementary table 2. Parameters used to compare models.**

| a_x | b_x | c_x | k_2x | k_3x | k_4x | a_y | b_y | c_y | k_2y | k_3y | k_4y | odel_nu |
|---|---|---|---|---|---|---|---|---|---|---|---|---|
| 0.01 | 1.75 | 0.75 | 0.5 | 0 | 0 | 0 | 0.1 | 0.05 | 0.5 | 0 | 0 | **A1** |
| 0.01 | 1.75 | 0.75 | 0.5 | 0 | 0 | 0 | 0.075 | 0.05 | 0.1 | 0 | 0 | **A2** |
| 0.01 | 1.75 | 0.75 | 0.85 | 0 | 0 | 0 | 0.1 | 0.05 | 0.5 | 0 | 0 | **A3** |
| 0.01 | 1.75 | 0.75 | 0.85 | 0 | 0 | 0 | 0.075 | 0.05 | 0.1 | 0 | 0 | **A4** |
| 0.01 | 1.75 | 0.75 | 0.5 | 0 | 0 | 0.17 | 0 | 0.05 | 0 | 0.01 | 0 | **A5** |
| 0.01 | 1.75 | 0.75 | 0.5 | 0 | 0 | 8.5 | 0 | 0.05 | 0 | 0.01 | 0 | **A6** |
| 0.01 | 1.75 | 0.75 | 0.85 | 0 | 0 | 0.17 | 0 | 0.05 | 0 | 0.01 | 0 | **A7** |
| 0.01 | 1.75 | 0.75 | 0.85 | 0 | 0 | 8.5 | 0 | 0.05 | 0 | 0.01 | 0 | **A8** |
| 0.01 | 1.38 | 0.75 | 0 | 0 | 0.05 | 0 | 0.1 | 0.05 | 0.5 | 0 | 0 | **A9** |
| 0.01 | 1.38 | 0.75 | 0 | 0 | 0.05 | 0 | 0.075 | 0.05 | 0.1 | 0 | 0 | **A10** |
| 0.01 | 1.472 | 0.75 | 0 | 0 | 0.05 | 0 | 0.1 | 0.05 | 0.5 | 0 | 0 | **A11** |
| 0.01 | 1.472 | 0.75 | 0 | 0 | 0.05 | 0 | 0.075 | 0.05 | 0.1 | 0 | 0 | **A12** |
| 0.01 | 1.38 | 0.75 | 0 | 0 | 0.05 | 0.17 | 0 | 0.05 | 0 | 0.01 | 0 | **A13** |
| 0.01 | 1.38 | 0.75 | 0 | 0 | 0.05 | 8.5 | 0 | 0.05 | 0 | 0.01 | 0 | **A14** |
| 0.01 | 1.472 | 0.75 | 0 | 0 | 0.05 | 0.17 | 0 | 0.05 | 0 | 0.01 | 0 | **A15** |
| 0.01 | 1.472 | 0.75 | 0 | 0 | 0.05 | 8.5 | 0 | 0.05 | 0 | 0.01 | 0 | **A16** |
| 0.05 | 5 | 0.75 | 0.5 | -1 | -1 | 0.005 | 0.005 | 0.1 | 0.05 | -1 | -1 | **B1** |
| 0.05 | 5 | 0.75 | 0.5 | -1 | -1 | 0.005 | 0.002 | 0.1 | 0.22 | -1 | -1 | **B2** |
| 0.05 | 3.26 | 0.75 | 0.25 | -1 | -1 | 0.005 | 0.005 | 0.1 | 0.05 | -1 | -1 | **B3** |
| 0.05 | 3.26 | 0.75 | 0.25 | -1 | -1 | 0.005 | 0.002 | 0.1 | 0.22 | -1 | -1 | **B4** |
| 0.05 | 5 | 0.75 | 0.5 | -1 | -1 | 0.005 | 0.005 | 0.1 | -1 | 0.3 | -1 | **B5** |
| 0.05 | 5 | 0.75 | 0.5 | -1 | -1 | 0.005 | 0.005 | 0.1 | -1 | 0.17 | -1 | **B6** |
| 0.05 | 3.26 | 0.75 | 0.25 | -1 | -1 | 0.005 | 0.005 | 0.1 | -1 | 0.3 | -1 | **B7** |
| 0.05 | 3.26 | 0.75 | 0.25 | -1 | -1 | 0.005 | 0.005 | 0.1 | -1 | 0.17 | -1 | **B8** |
| 0.05 | 2.5 | 0.75 | -1 | -1 | 0.4 | 0.005 | 0.005 | 0.1 | 0.05 | -1 | -1 | **B9** |
| 0.05 | 2.5 | 0.75 | -1 | -1 | 0.4 | 0.005 | 0.002 | 0.1 | 0.22 | -1 | -1 | **B10** |
| 0.06 | 2.1 | 0.75 | -1 | -1 | 0.8 | 0.005 | 0.005 | 0.1 | 0.05 | -1 | -1 | **B11** |
| 0.06 | 2.1 | 0.75 | -1 | -1 | 0.8 | 0.005 | 0.002 | 0.1 | 0.22 | -1 | -1 | **B12** |
| 0.05 | 2.5 | 0.75 | -1 | -1 | 0.4 | 0.005 | 0.005 | 0.1 | -1 | 0.3 | -1 | **B13** |
| 0.05 | 2.5 | 0.75 | -1 | -1 | 0.4 | 0.005 | 0.005 | 0.1 | -1 | 0.17 | -1 | **B14** |
| 0.06 | 2.1 | 0.75 | -1 | -1 | 0.8 | 0.005 | 0.005 | 0.1 | -1 | 0.3 | -1 | **B15** |
| 0.06 | 2.1 | 0.75 | -1 | -1 | 0.8 | 0.005 | 0.005 | 0.1 | -1 | 0.17 | -1 | **B16** |

**Supplementary table 3. Best fit parameters for pro inflammatory-IL10 circuit.**

| patient | p1 | p2 | p3 | p4 | p5 | p6 | p7 | p8 | r2 | weighted r2 | redictiv |
|---|---|---|---|---|---|---|---|---|---|---|---|
| 4 | 1.13 (0.65, 2.36) | 4.09 (3.71, 4.35) | 0.77 (0.69, 0.84) | 1.32 (0.54, 3.02) | 0.42 (0.07, 0.81) | 0.02 (-0.10, 0.10) | 0.00 (-0.08, 0.03) | 0.27 (0.01, 0.85) | 0.1666 | 0.4716 | 92.50% |
| 5 | 4.43 (0.22, 5.02) | 9.86 (1.87, 11.91) | 6.84 (2.82, 25.26) | 3.57 (0.52, 4.72) | 2.06 (0.36, 5.17) | 0.03 (0.02, 8.85) | 0.01 (0.00, 14.05) | 1.21 (0.58, 9.76) | 0.1401 | 0.6686 | 40.00% |
| 6 | 1.61 (1.38, 2.00) | 6.13 (5.80, 6.44) | 1.01 (0.97, 1.05) | 1.94 (1.59, 2.53) | 0.43 (0.04, 0.94) | 0.15 (0.14, 0.16) | 0.28 (0.25, 0.31) | 0.60 (0.07, 1.04) | 0.8350 | 0.8510 | 20.00% |
| 7 | 6.08 (2.24, 11.69) | 4.72 (2.19, 9.69) | 1.69 (1.31, 2.92) | 7.44 (2.01, 17.70) | 2.67 (0.16, 6.58) | 0.05 (-0.13, 0.06) | 0.00 (-0.03, 0.02) | 0.71 (0.11, 1.21) | -7.0152 | 0.7243 | 72.22% |
| 9 | 5.17 (4.17, 5.87) | 4.26 (4.02, 4.67) | 1.63 (1.19, 1.90) | 3.76 (3.12, 4.97) | 0.80 (0.09, 1.78) | 0.27 (0.22, 0.28) | 0.35 (0.25, 0.38) | 0.18 (0.02, 0.49) | 0.7130 | 0.8178 | 37.50% |
| 10 | 3.10 (1.67, 5.48) | 5.55 (4.05, 8.40) | 5.45 (4.65, 6.86) | 1.82 (0.35, 4.53) | 0.47 (0.04, 1.24) | 0.18 (0.15, 0.21) | 0.23 (0.18, 0.28) | 0.61 (0.11, 1.58) | 0.8506 | 0.9130 | 28.95% |
| 11 | 1.49 (0.71, 3.08) | 3.21 (2.54, 3.58) | 0.91 (0.83, 1.24) | 0.80 (-0.07, 2.56) | 0.64 (0.15, 1.04) | 0.15 (0.09, 0.17) | 0.07 (-0.10, 0.09) | 0.24 (0.03, 0.82) | 0.6687 | 0.8045 | 90.00% |
| 12 | 3.91 (1.12, 7.60) | 1.94 (1.26, 2.29) | 1.97 (0.81, 2.50) | 0.89 (0.09, 2.49) | 0.01 (0.00, 0.05) | 0.37 (0.13, 0.56) | 0.45 (0.10, 0.74) | 0.06 (0.00, 0.25) | 0.7240 | 0.8125 | 82.50% |
| 13 | 4.45 (3.59, 5.17) | 4.99 (4.80, 5.32) | 1.88 (1.77, 2.01) | 4.86 (3.74, 5.89) | 1.94 (1.06, 2.59) | 0.30 (0.25, 0.36) | 0.53 (0.42, 0.66) | 0.18 (0.01, 1.17) | 0.8314 | 0.8750 | 25.00% |
| 14 | 11.92 (1.88, 19.97) | 3.73 (3.62, 4.32) | 1.50 (0.85, 2.11) | 9.40 (0.93, 16.23) | 0.29 (0.04, 2.57) | 0.58 (0.42, 0.61) | 0.40 (0.23, 0.43) | 1.74 (0.32, 3.13) | 0.0932 | 0.6592 | 15.00% |
| 15 | 10.85 (1.08, 13.28) | 1.43 (0.33, 1.85) | 5.23 (1.14, 24.63) | 4.68 (2.45, 47.26) | 0.53 (0.06, 14.03) | 0.33 (0.07, 6.84) | 0.64 (0.11, 15.09) | 0.53 (0.01, 19.86) | 0.5443 | 0.8455 | 66.67% |
| 16 | 1.10 (0.47, 2.94) | 2.39 (1.91, 3.74) | 2.64 (2.42, 2.94) | 0.53 (0.04, 2.13) | 0.24 (0.02, 0.63) | 0.09 (0.07, 0.11) | 0.17 (0.11, 0.23) | 0.21 (0.02, 0.59) | 0.8650 | 0.7850 | 27.50% |
| 17 | 2.41 (0.75, 4.07) | 7.34 (5.90, 9.31) | 5.23 (3.28, 7.52) | 1.63 (0.03, 3.53) | 1.09 (0.72, 1.66) | 0.19 (0.14, 0.27) | 0.39 (0.23, 0.58) | 0.18 (0.02, 0.83) | 0.7910 | 0.9243 | 32.50% |

| | | | | | | | | | | | |
|---|---|---|---|---|---|---|---|---|---|---|---|
| 18 | 6.12 (5.96, 6.39) | 8.39 (8.19, 9.70) | 1.50 (1.43, 2.01) | 3.31 (2.42, 3.43) | 1.07 (0.24, 1.15) | 7.48 (5.70, 7.78) | 3.67 (2.77, 3.86) | 0.04 (0.00, 0.82) | -0.8631 | 0.9579 | 53.12% |
| 1 | 3.19 (2.99, 3.64) | 3.34 (3.00, 3.62) | 1.71 (1.42, 2.08) | 2.96 (2.66, 3.77) | 0.21 (-0.14, 0.99) | 0.01 (-0.14, 0.03) | 0.06 (0.00, 0.22) | 0.67 (0.06, 0.85) | -1.5206 | 0.7132 | 72.50% |
| 2 | 0.41 (0.29, 0.59) | 6.98 (5.12, 10.71) | 1.26 (1.17, 1.40) | 0.08 (0.01, 0.59) | 0.24 (0.02, 0.57) | 0.10 (0.08, 0.13) | 0.20 (0.14, 0.28) | 0.21 (0.02, 0.57) | 0.7156 | 0.6371 | 47.50% |
| 3 | 4.66 (4.05, 5.98) | 5.98 (5.08, 6.27) | 4.83 (3.79, 5.19) | 3.90 (2.35, 5.68) | 0.27 (0.04, 1.53) | 0.33 (0.26, 0.37) | 0.71 (0.47, 0.83) | 1.45 (0.58, 2.20) | 0.9357 | 0.9720 | 50.00% |
| 7 | 2.37 (1.50, 6.53) | 2.64 (2.00, 7.09) | 1.55 (1.25, 2.03) | 2.17 (1.75, 9.06) | 0.34 (0.02, 1.63) | 0.05 (-0.02, 0.06) | 0.00 (-0.14, 0.02) | 1.01 (0.26, 3.04) | 0.7880 | 0.7153 | 83.33% |

**Supplementary table 4. Parameter range for sensitivity analysis**

| | a | b | c | d | f | g |
|---|---|---|---|---|---|---|
| **Default value** | 1 | 1 | 100 | 2 | 1 | 1 |
| **Threshold for activation** | 0.1 - 10 | - | 20-500 | 0.01 - 20 | - | - |
| **Response amplitude** | 0.5 - 2.5 | 0.1 - 5 | 20 - 500 | 0.01 - 10 | 0.1 - 2.5 | 0.1 - 10 |
| **Refractory Period** | 0.5 - 2.5 | 0.1 - 5 | 20 - 500 | 0.01 - 10 | 2.5 - 10 | 0.1 - 2 |

**Supplementary Figure 1. Type I interferon pulse ends before viral load decreases in mice infected with IAV and RSV.**

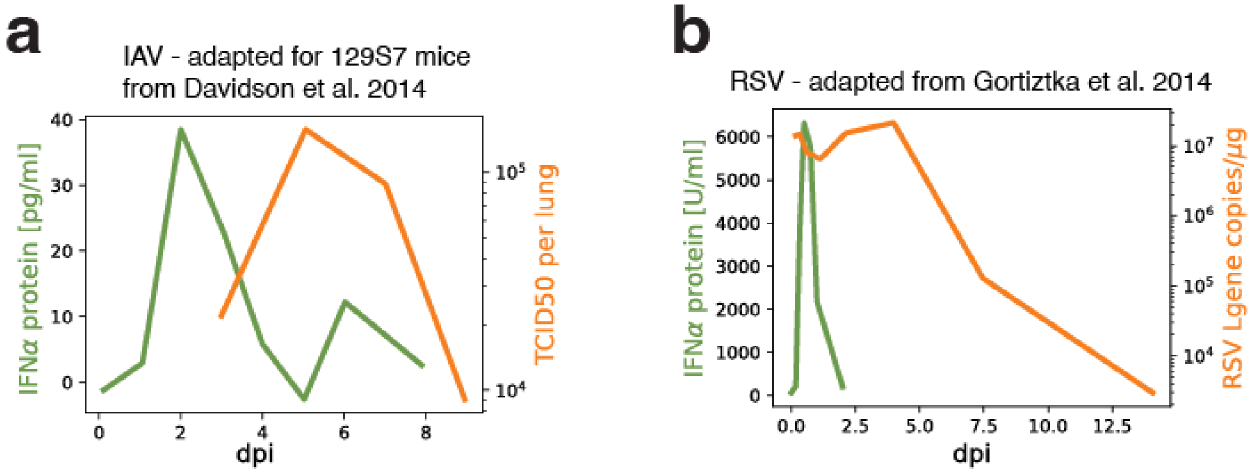


a) Type I interferon levels (green) vs. viral load (orange) measured in 129S7 mice infected with IAV over the first 8 days post-infection (dpi). Adapted from Davidson et al. 2014.

b) Type I interferon levels (green) vs. viral load (orange) measured in mice infected with RSV over the first 14 days post-infection. Adapted from Goritzka et al. 2014.

**Supplementary figure 2. All excitable cytokine circuit topologies.** This list is corresponding with supplementary table 1.

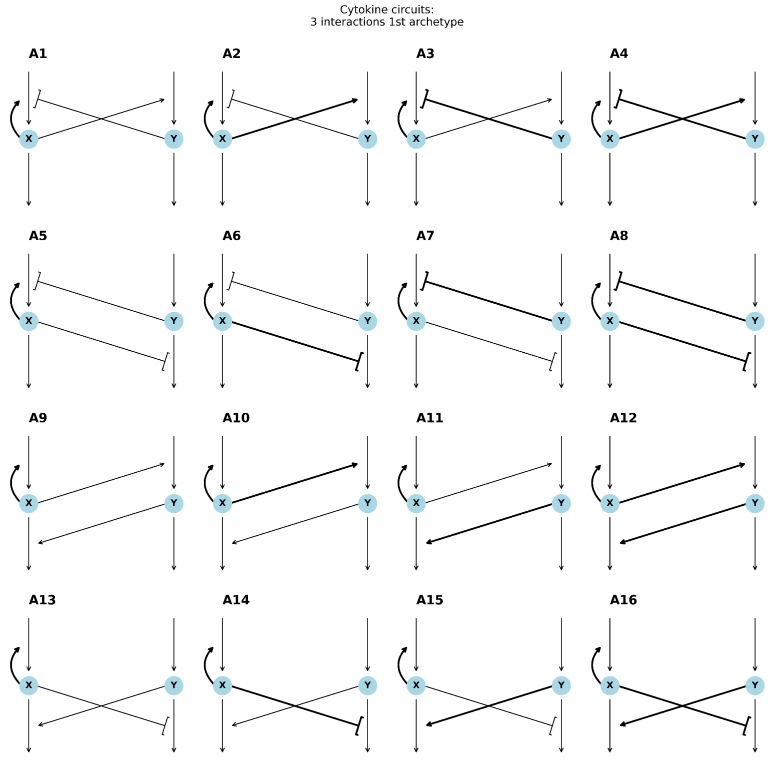


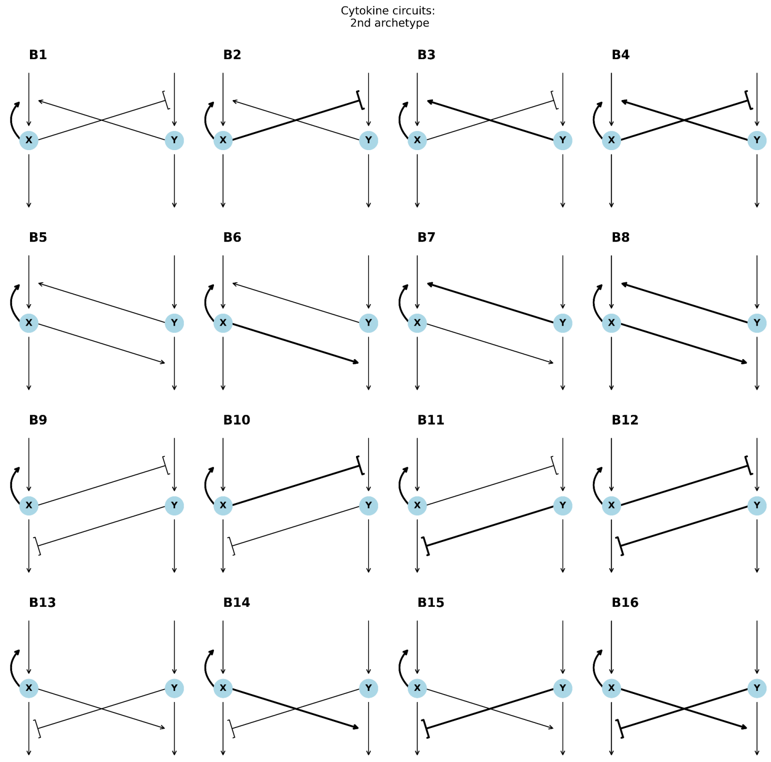

### Supplementary figure 3. Pareto fronts of models in all metrics

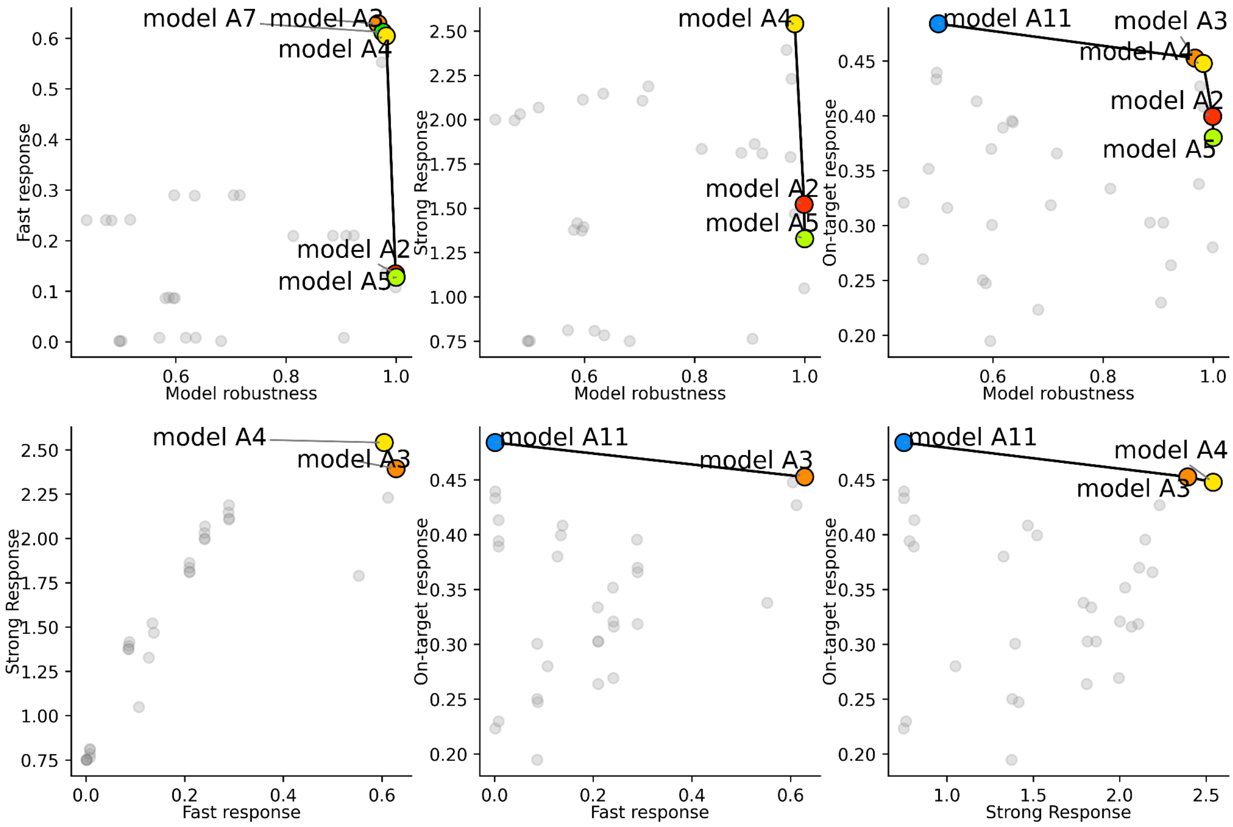

### Supplementary Figure 4. Pro inflammatory cytokines -IL10 model fit to all patients.

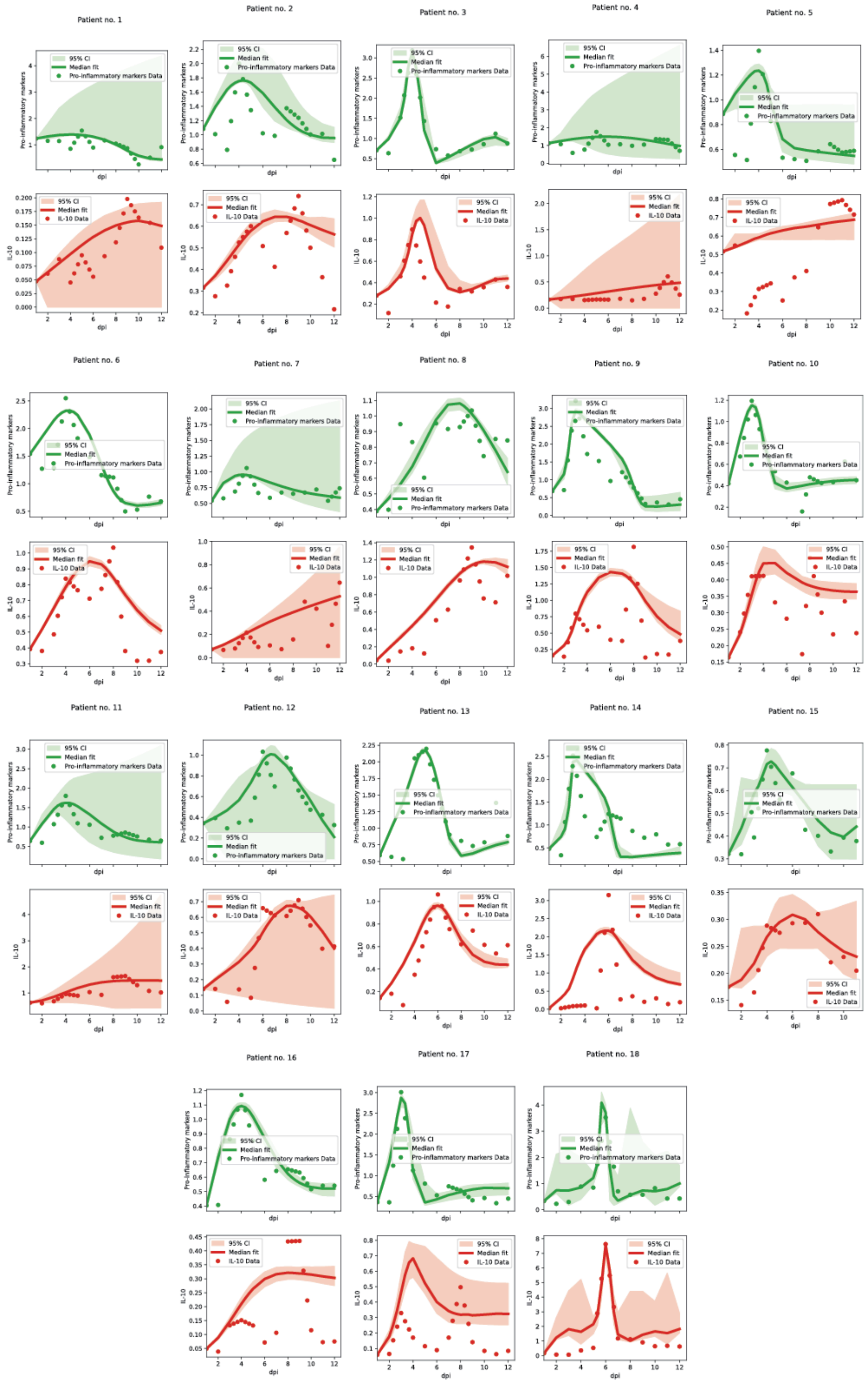

**Supplementary Figure 5. Motifs distribution in the cytokine network.** X- axis is the z score of each motif. Blue bars are significantly enriched (z-score>3). The red bar is the excitable motif.

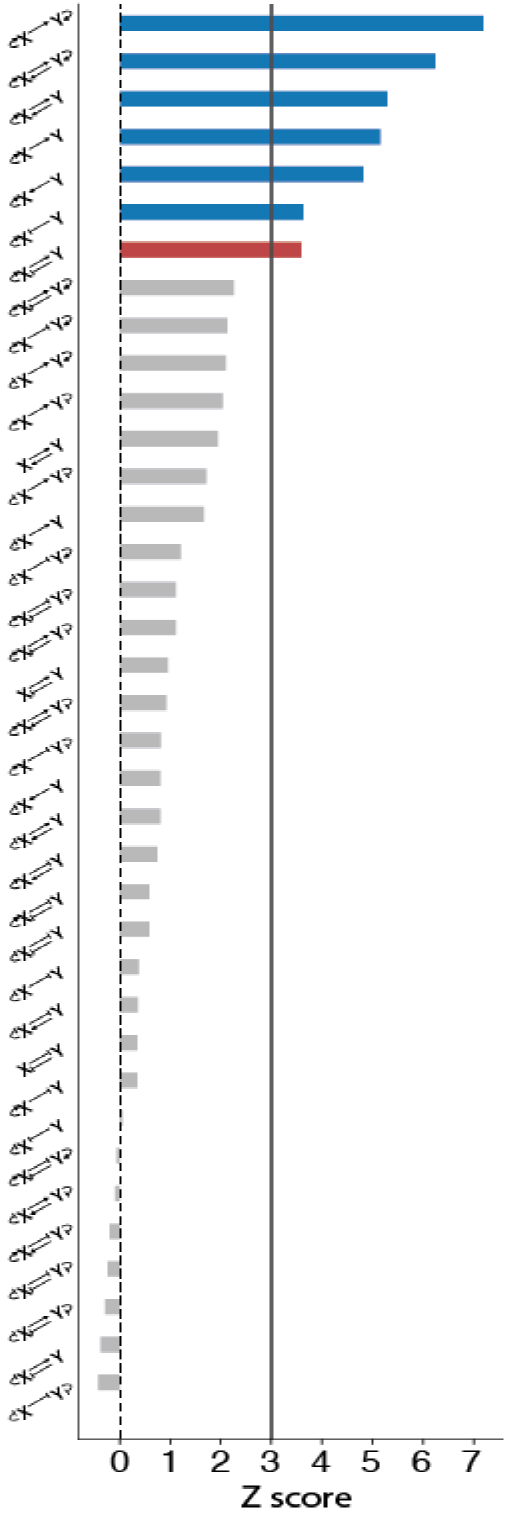
